# Calibrating seed-based alignment heuristics with Sesame

**DOI:** 10.1101/619155

**Authors:** Guillaume J. Filion, Ruggero Cortini, Eduard Zorita

## Abstract

The increasing throughput of DNA sequencing technologies creates a need for faster algorithms. The fate of most reads is to be mapped to a reference sequence, typically a genome. Modern mappers rely on heuristics to gain speed at a reasonable cost for accuracy. In the seeding heuristic, short matches between the reads and the genome are used to narrow the search to a set of candidate locations. Several seeding variants used in modern mappers show good empirical performance but they are difficult to calibrate or to optimize for lack of theoretical results. Here we develop a theory to estimate the probability that the correct location of a read is filtered out during seeding, resulting in mapping errors. We describe the properties of simple exact seeds, skip-seeds and MEM seeds (Maximal Exact Match seeds). The main innovation of this work is to use concepts from analytic combinatorics to represent reads as abstract sequences, and to specify their generative function to estimate the probabilities of interest. We provide several algorithms, which combined together give a workable solution for the problem of calibrating seeding heuristics for short reads. We also provide a C implementation of these algorithms in a library called Sesame. These results can improve current mapping algorithms and lay the foundation of a general strategy to tackle sequence alignment problems. The Sesame library is open source and available for download at https://github.com/gui11aume/sesame.

## 1 Introduction

### 1.1 Mapping reads to genomes

Suppose that we use and imperfect instrument to sequence a DNA fragment drawn at random from a known genome; how can we identify the location of the fragment in the genome?

We will refer to this question as the *true location problem*. It has applications in countless domains, like discovering disease-causing mutations; diagnosing viral, bacterial or fungal infections; diagnosing cancer and choosing appropriate therapies; selecting breeds of interest in agriculture; tracing human migrations from ancient bones; diagnosing genetic diseases at birth; finding compatible graft donors *etc*.

At first sight, whether a read can be mapped to its correct location seems to depend only on its length and on the error rate of the sequencer. However, there is another important factor to consider: most genomes contain repeated sequences, *i.e.* relatively large subsequences that are present at more than one location. For instance, we cannot map the fragment with certainty if there exists an exact copy of the sequence somewhere else in the genome, because we cannot know which of the two copies was sequenced.

This case is rare though. Repeated sequences are in general not identical but merely homologous — meaning that their similarity is unlikely to occur by chance. So when the DNA fragment has one or more duplicates in the genome, we can still map it, as long as we can distinguish the duplicates. This in turn depends on their similarity with the target fragment and on the error rate of the sequencer.

Since repeats play a central role in the problem at hand, we give the terms “targets” and “duplicates” a meaning that will facilitate the exposition of the theory.

#### Definition 1.

*The target is the DNA fragment that was sequenced. Duplicates are sequences of the genome that are homologous to the target*.

Because sequencing errors can convert the sequence of the target into one of its duplicates, the *true* location of a DNA fragment is not always the *best* candidate (as measured by the identity between the sequence and the candidate location). This leads to the following question: how can we identify the *best* location of the fragment in the genome?

We will refer to this task as the *best location problem*. It amounts to finding the optimal alignment between two sequences, and for this reason has received substantial attention in bioinformatics. For simplicity we will assume that the true location is always the best, and we will often confound these two ideas.

### 1.2 Seeding heuristics

There exist algorithms to solve the best location problem [1, 2], but they are too slow to process the large amount of data generated by modern sequencers. Instead, one uses heuristic methods, *i.e.* algorithms that run faster, but that may return an incorrect result [3].

The most popular heuristic for the best location problem is a filtration method called “seeding”. The principle is to first identify short regions of high similiarity between the read and the genome, and then run an exact alignment algorithm at the candidate locations. This is for instance the general strategy of the popular alignment algorithm BLAST [4].

If seeding is fast, we can quickly zoom in on a small set of candidate locations and reduce the input size of the (slow) alignment algorithm. The disadvantage is that the target location is not always detected and included in the candidate set. When that happens, the read cannot be mapped to the true location because it has been filtered out at the seeding step.

The slowness of exact alignment algorithms imposes a trade-off between speed and accuracy: If the seeding step is set to produce many candidates, it is likely to retrieve the true location. The downside is that processing all the candidates would take long. Conversely, if the seeding step is set to produce few candidates, the process would run faster but the target is less likely to be in the candidate set.

Importantly, the trade-off depends on the seeding method. This means that we can develop faster mapping algorithms at no cost for accuracy, as long as we can find better seeding strategies. Progress on this line of research has largely benefited from the improvement of computer hardware and from the development of indexing algorithms. Literature on this topic is already abundant. Some examples of seeding algorithms are given in references [5–11]. References [12] and [13] present high-level comparisons of different designs, and reference [14] gives a global overview of filtration methods in pattern matching.

### 1.3 The two types of seeding failure

Filtering methods are considered to fail whenever the target is not in the candidate set. For seeding heuristics, we need to distinguish two different kinds of failure. In the first kind, the candidate set contains a duplicate of the target but does not contain the target itself; in the second kind, the candidate set contains neither.

The distinction is important because duplicates of the target are similar to the read (due to their similarity to the target), so a failure of the first kind is difficult to distinguish from a success. A failure of the second kind is easier to recognize because in this case the best hit is not similar to the read, as we will highlight in section 7.

Before going further, we introduce some nomenclature that will be useful to simplify the exposition of the theory.

#### Definition 2.

*The seeding process is said to be*

i. “on target” if the candidate set contains the target,
ii. “off target” if the candidate set contains a duplicate but not the target,
iii. “null” if the candidate set contains neither.

If a seed-based mapping algorithm returns an incorrect location, then it must be the case that the best location was never tested in the post-filtration phase (otherwise it would have been found by the exact alignment algorithm).

This observation is fundamental: in order to estimate the error rate of a mapping algorithm, we must know how often the associated seeding heuristic is off target. For lack of theoretical framework, this information is missing, meaning that current mappers are not properly calibrated.

Our focus here is to develop a theory to estimate the probability that seeding is off target, when using the most popular seeding heuristics for short reads. Previous work pioneered a method to compute seeding probabilities but it did not distinguish off-target from null seeding [15,16], and therefore did not provide a way to estimate the frequency of mapping errors. Other authors investigated the reliability of mapping algorithms [17], but they focused on the probability that random sequences have a single hit, recognizing that solving the problem requires taking into consideration the repeat structure of the genome. Here we address this problem in general terms and we provide a practical way to estimate the mismapping rate in complex genomes.

The rest of this document is not organized by logical units but by increasing level of difficulty. The order of the examples is meant to ease the acquisition of the main concepts, meaning that we sometimes go back and forth between different types of seeding strategies. We first give some background on the different types of seeding methods considered here; we then introduce the computational strategy based on transfer matrices; finally we delve into each specific seeding method.

## 2 Seeds

The term “seed” changes meaning with different authors. It can designate a part of the read, a part of the genome, a particular sequence motif or a structured pattern of matches. Also, the term does not always imply an exact match. For instance, the algorithm PatternHunter [6] uses “spaced seeds” that tolerate mismatches. To avoid potential confusion, we will adopt the following convention:

### Definition 3.

*A seed is a subsequence of the read that has size at least γ (defined by the context of the problem) and that has at least one perfect match in the reference genome*.

When a seed matches a particular location of the genome, we say that it is a “seed for that location”. For instance, we often use expressions such as “a seed for the target” or “a seed for a duplicate”.

This definition presents a computational challenge: to know if a given subsequence is a seed we need to know if it exists somewhere in the genome. This is a non trivial problem in itself, but fortunately we can use practical methods to solve it, even when the reference genome is very large.

These algorithms are crucial for the present theory, but describing them in depth is outside the scope of this document. Let us just mention that all the methods belong to a family known as exact offline string matching algorithms, where “offline” means that sequences are looked up in an index instead of the genome itself. Online methods can be used when the reference genome is not indexed [18], but this case is of little relevance in the present context.

The index is usually a hash table or a variant of the so-called FM-index [20, 21]. Hash tables are typically used to index *k*-mers, which makes them useful to search seeds of fixed size *k* (see [19] for a recent benchmark of *k*-mer hashing algorithms). In contrast, some text-indexing structures have no set size so they can be used to search seeds of different lengths. For instance, the FM-index is a compact data structure based on the suffix array [22] and on the Burrows-Wheeler transform [23], emulating a suffix trie with a much smaller memory footprint [20, 21].

Other methods can be efficient (*e.g.*, running a bisection on the suffix array [24]) but the FM-index is currently the most popular choice for seeding methods. For MEM seeds, it is even the only practical option [25–28]. However, the detail is of little interest here: we simply assume that seeds are known at all times without ambiguity because this problem has several practical solutions.

### 2.1 Exact seeds

Exact seeds have a fixed size *γ*. When using exact seeds, all the perfect matches of size *γ* are collected to constitute the candidate set. This seeding heuristic was used in the first version of BLASTN [4], but is no longer used because it produces many overlapping matches within the same regions.

Fig. 1 shows a read produced by an imperfect sequencer. The sequenced DNA fragment comes from a region of the genome that has three duplicates and the read has three miscalled nucleotides (indicated by stars).

**Figure 1:**
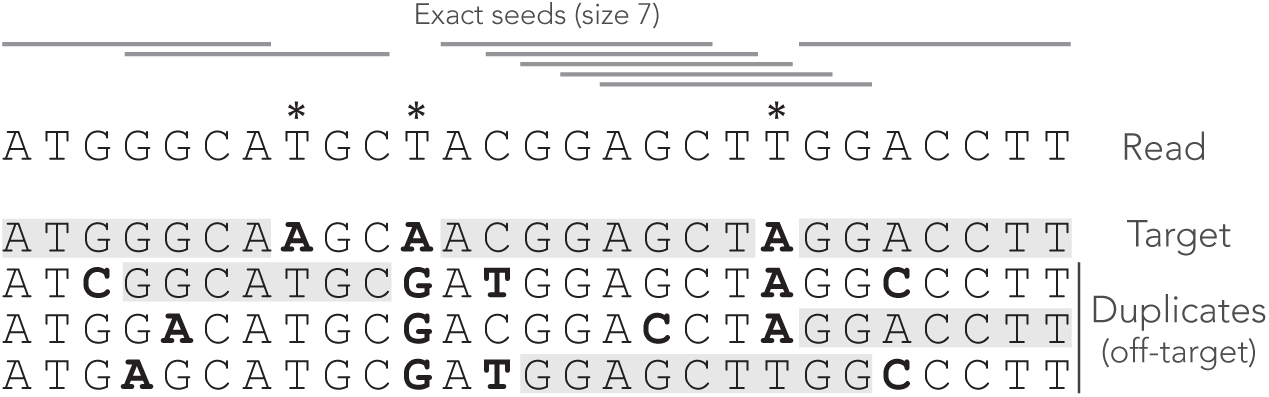
Example of exact seeds. A sequenced DNA fragment (Read) is shown above the actual molecule (Target), and above homologous sequences also present in the genome (Duplicates). Sequencing errors are indicated by a star (each marked T was actually an A that was misread). The mismatches between the genomic sequences and the read are indicated in bold. The exact seeds of size *γ* = 7 are indicated as horizontal grey lines above the read. The matching regions in the genomic sequences are shadowed in grey. Several overlapping seeds accumulate at the center of the read, which is typical for this seeding method.

Observe that erroneous nucleotides can belong to exact seeds because they sometimes match one of the duplicates. For instance, the first sequencing error matches all the duplicates and belongs to an off-target seed. However, sequencing errors that are mismatches for *all* the sequences cannot belong to a seed. This is the case of the second sequencing error in this example, where there is a local deficit of seeds.

The middle of the read shows some clutter where consecutive seeds match consecutive sequences at the same location. This is typical for exact seeds and constitutes a nuisance for the implementation. Indeed, it is a waste of computer resources to discover matches in sequences that are already in the candidate set. In addition, this seeding method is not particularly sensitive compared to spaced seeds [6] so it is used only in few specific applications. Nevertheless, it will be extremely useful for the development of the present theory.

### 2.2 Skip seeds

Skip seeds have a fixed size *γ*, but unlike exact seeds they cannot start at every nucleotide. Instead, a certain amount of nucleotides is skipped between every seed. This is a way to reduce the overlapping matches at the same location, at the cost of sensitivity. This seeding heuristic is the core of Bowtie2 [29], where seeds have size *γ* = 16 and are separated by 10 nucleotides (9 positions are skipped). We will refer to seeds where *n* nucleotides are skipped as “skip-*n* seeds”. For instance, Bowtie2 uses skip-9 seeds.

Fig. 2 shows what happens when exact seeds are replaced by skip-1 seeds on the read of Fig. 1. Here the size is still *γ* =7 but 1 nucleotide is skipped between seeds. This amounts to removing every second seed. The consequence is that there are fewer overlapping matches at the center of the read, but the only seed for the second duplicate is lost. This is a rather positive outcome because there is one off-target location fewer in the candidate set, but the same might happen to the target.

**Figure 2:**
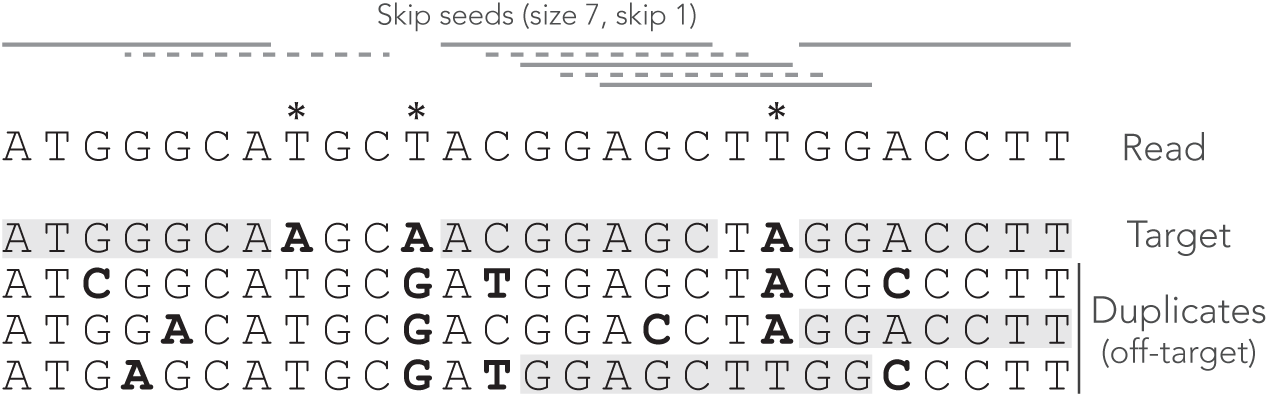
Example of skip seeds. The sequences and the annotations are the same as in Fig. 1, but here we use skip-1 seeds. In other words, seeds can never start at nucleotides 2, 4, 6 *etc*. To highlight the difference with Fig. 1, the missing seeds are represented by dotted lines.

It is clear that skipping nucleotides reduces the sensitivity of the seeding step, but to what extent? One could test this empirically, but the answer depends on the seed length, the number of nucleotides that are skipped, the error rate of the sequencer and the size of the reads. The present theory will allow us to make general statements about the performance of skip seeds against exact seeds in different contexts.

### 2.3 MEM seeds

MEM seeds (where MEM stands for Maximal Exact Match) are somewhat harder to define. Unlike exact seeds and skip seeds, their size is variable. They are used in BWA-MEM [30] where they give good empirical results. To describe MEM seeds, let us first clarify the meaning of “Maximal Exact Match”.

#### Definition 4.

*A Maximal Exact Match (MEM) is a subsequence of the read that is present in the reference genome and that cannot be extended — either because the read ends or because the extended subsequence is not in the genome*.

A *MEM seed* is simply a MEM of size size *γ* or greater. Fig. 3 shows what happens when using MEM seeds on the read of Fig. 1. Observe that the clutter at the center of the read has disappeared because consecutive matches are fused into a few MEM seeds.

Two consecutive MEM seeds can overlap, in which case they always match distinct sequences of the genome (otherwise neither of them would be a MEM seed). There does not have to be any overlap though, because a nucleotide can be a mismatch against *all* the sequences, like the second read error for instance. Note that a MEM does not always match a single sequence of the genome.

For instance, the rightmost MEM seed matches two distinct genomic sequences. This case motivates the following definition, which will play an important role later.

**Figure 3:**
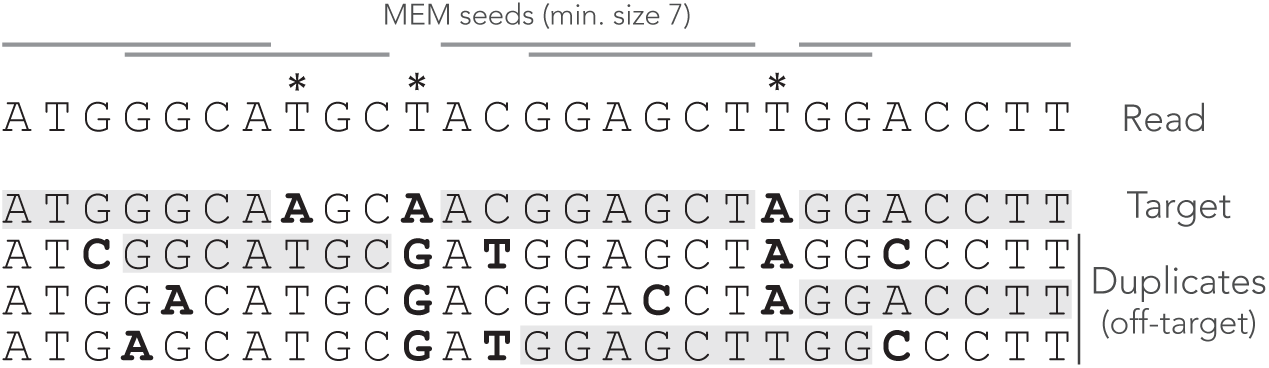
Example of MEM seeds. The sequences and the annotations are the same as in Fig. 1, but here we use MEM seeds of minimum size *γ* = 7. The clutter at the center of the read has disappeared and there is at least one seed for each sequence.

#### Definition 5.

*A* strict *MEM seed has a single match in the genome. A* shared *MEM seed has several matches in the genome*.

Compared to seeds of fixed size, MEM seeds have two counter-intuitive properties. The first is that there are cases where there cannot be any on-target seed, even when changing the sensitivity parameter *γ*. Fig. 4 shows such an example. Even though there is a single sequencing error, the read has no MEM seed for the target. Lowering *γ* does not change this, so there is no way to discover the true location using MEM seeds (even though it is the best location).

The second counter-intuitive property is that shortening a read can sometimes generate a MEM seed for the target. Fig. 5 shows an example of this case. There is no MEM seed for the target, but there would be if the read were two nucleotides shorter on the right side. Indeed, in this case there would be a shared MEM seed matching the the target and the first duplicate (provided *γ ≤* 12).

These examples show that MEM seeds can perform worse than seeds of fixed size. MEM seeds yield fewer candidates and therefore speed up the mapping process, but the question is at what cost? The theory developed here will allow us to compute the probability that a read has no MEM seed for the target and thus that the true location is missed at the seeding stage.

**Figure 4:**
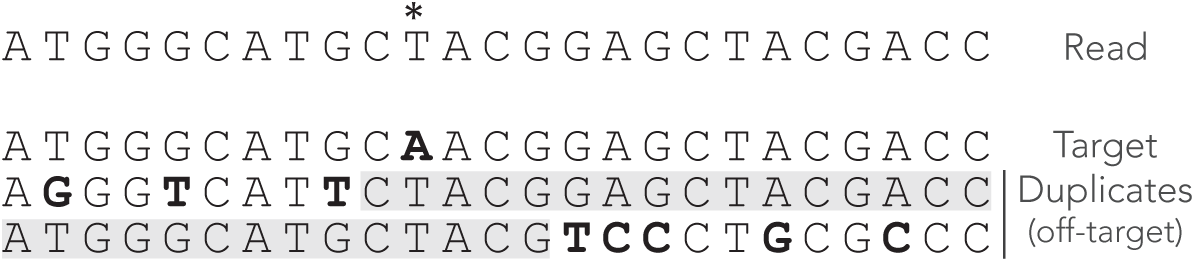
Issues with MEM seeds: inaccessible targets. The read, the MEM seeds and the sequences are represented as in Fig. 3. The MEM seeds matching the two duplicates at the bottom effectively hide the target, so it cannot be discovered. This can occur even when the true location is the best candidate and when there is a single error on the read.

**Figure 5:**
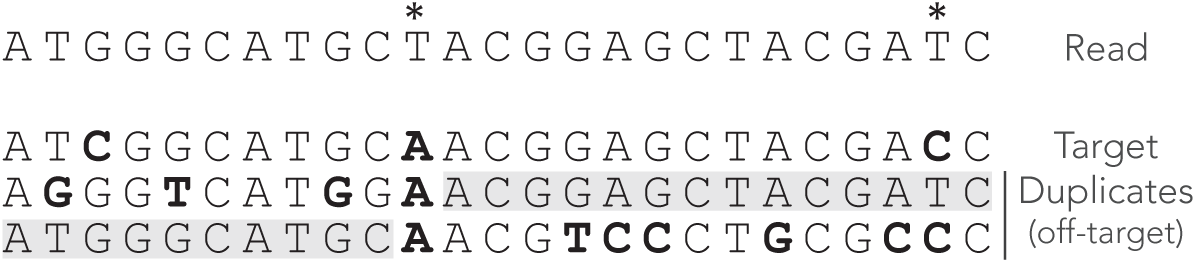
Issues with MEM seeds: too long reads. The read, the MEM seeds and the sequences are represented as in Fig. 3. There would be a good seed (shared) if the read were two nucleotides shorter. The true location is hidden by the last two nucleotides.

## 3 Model and strategy

### 3.1 Sequencing errors and divergence of the duplicates

We now need to model the sequencing and duplication processes so that we can compute the probabilities of the events of interest. We assume that the sequencing instrument has a constant substitution rate *p*, and that insertions and deletions never occur. When a substitution occurs, we assume that the instrument is equally likely to output any one of the remaining three nucleotides. This corresponds more or less to the error spectrum of the Illumina sequencing technology [31].

Next, we assume that the target sequence has *N*≥0 duplicates, so that there are *N* off-target sequences. We further assume that duplication happened instantaneously at some point in the past and that all *N* +1 sequences diverge independently of each other at a constant rate. In other words, we ignore the complications due to the genealogy of the duplication events. Instead, we simply assume that at each nucleotide position, any given duplicate is identical to the target with probability 1−*µ*. If it is not, we assume that the duplicate sequence can be any of one the three remaining nucleotides (*i.e.* each is found with probability *µ/*3).

Note that read errors are always mismatches against the target (because we assume that the target is the true sequence), and they match each duplicate with probability *µ/*3. Correct nucleotides are always matches for the target, and they match each duplicate with probability 1 − *µ*.

Before going further, we also need to move a practical consideration out of the way. Seeds can match any sequence of the genome, not just the target or the duplicates. However, we will ignore matches in the rest of the genome because such random matches are unlikely to cause a mapping error when seeding is off target, contrary to matches in duplicates. Neglecting those will greatly simplify the exposition of the theory without loss of generality. We will explain in section 7 how to deal with this practical case and and how to identify those matches as off target. Until then, we will consider that the target and the duplicates are the only sequences in the reference genome.

### 3.2 Weighted generating functions

The central object of analytic combinatorics is the generating function, and for our purpose we will use a special kind known as *weighted generating function*.

#### Definition 6.

*Let* 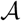 *be a set of combinatorial objects such that* 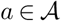 *has a size* 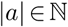 *and a weight w*(*a*) ℝ^+^. *The weighted generating function of* 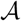 *is defined as*

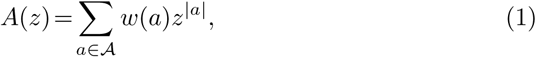

*Expression (1) also defines a sequence* (*a*_*k*_)_*k≥*0_ *such that*

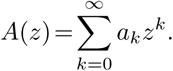

*By definition* 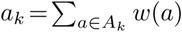, *where A*_*k*_ *is the class of objects of size k in* 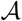. *The number a*_*k*_ *is the total weight of objects of size k*.

To give an example, assume that a particular symbol, say ⇓, has a probability of occurence equal to *p*. The weighted generating function words containing only this symbol is *pz*. The weight of the word is its probability (here equal to *p*) and the size is its length in number of symbols (here 1).

In this document we will focus on the weighted generating function *A*(*z*) of the set 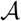 of reads that do not have on-target seeds (*i.e.* reads for which seeding is either null or off target). The weight of a read is its probability of occurrence and the size *k* is its number of nucleotides. The coefficient *a*_*k*_ is thus the proportion of reads of size *k* that do not have an on-target seed, which is the quantity of interest.

The motivation for introducing weighted generating functions is that operations on combinatorial objects translate into operations on their weighted generating functions. If *A*(*z*) and *B*(*z*) are the weighted generating functions of two mutually exclusive sets 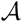 and 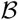, the weighted generating function of 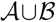 is *A*(*z*)+*B*(*z*), as evident from expression (1). Size and weight can be defined for pairs of objects in 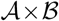 as (*a, b*) = *a* + *b* and *w*(*a, b*)=*w*(*a*)*w*(*b*). In other words the sizes are added and the weights are multiplied. With this convention, the weighted generating function of the Cartesian product 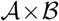 is *A*(*z*)*B*(*z*). This simply follows from expression (1) and from

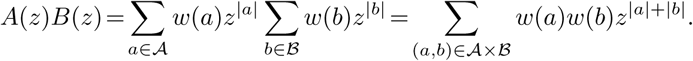

These two operations are all we need in order to compute the weighted generating functions of the reads of interest. Addition corresponds to creating a new family by merging reads from families 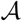 and 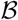. Multiplication corresponds to creating a new family by concatenating reads from families 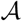 and 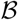.

### 3.3 Analytic representation

The analytic combinatorics framework relies on a strategy referred to as the *symbolic method* [33]. The idea is to combine simple objects into more complex objects. Each combinatorial operation on the objects corresponds to a mathematical operation on their weighted generating functions. One can thus obtain the weighted generating function of complex objects, whose coefficients *a*_*k*_ are the probabilities of interest.

As explained in [15] and [16], we recode the reads in alphabets of custom symbols and we specify a construction plan of the reads using a *transfer matrix M* (*z*). The transfer matrix specifies which types of segments can follow each other in the reads of interest: the entry at coordinates (*i, j*) is the weighted generating function of segments of type *j* that can be appended to segments of type *i*.

*M* (*z*) contains the weighted generating functions of all the reads that consist of a single fragment. From the basic operations on weighted generating functions, *M* (*z*)^*s*^ contains the weighted generating functions of all the reads that consist of *s* segments. Therefore, the entry at coordinates (*i, j*) of the matrix *M* (*z*)+ *M* (*z*)^2^ + *M* (*z*)^3^ + … = *M* (*z*) ⋅ (*I −M* (*z*))^*−*1^ is the weighted generating function of the reads of any size and any number of segments, that end with a segment of type *j* and that can be appended to a segment of type *i*. The examples below will clarify the key steps of this strategy.

A complete description of how to compute seeding probabilities with the symbolic method can be found in [15, 16]. The interested readers can also find more about analytic combinatorics in the popular textbooks [32, 33].

### 3.4 Example 1: on-target exact seeds

We highlight the strategy above with an example that will turn out to be central for the development of the theory. In addition, it is simple enough to provide a gentle introduction to the general methodology. This example was described in detail in [15] and in [16], but we repeat it here with a different formalism to fit the present manuscript.

The first step is to note that the nucleotide sequence of the read is irrelevant. Indeed, the read has an on-target seed if and only if it contains a stretch of *γ* nucleotides without error. For this reason, we recode reads as sequences of correct or erroneous nucleotides.

We define the mismatch alphabet 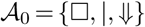, where □ represents a correct nucleotide, ⇓ represents an erroneous nucleotide and is special symbol appended after the last nucleotide to mark the end of the read. It is not associated to a nucleotide and therefore has size 0.

This recoding allows us to partition the read in an important way.

#### Definition 7.

*Every symbol of the alphabet that is different from* □ *is called a terminator. A segment is a terminator preceded by 0 or more symbols* □. *The last segment of the read is terminated by the symbol | and is called the tail*.

Since the decomposition of the read in segments is unique, we can view a read as a sequence of segments with a tail, instead of a sequence of nucleotides. Fig. 6 shows an example of decomposition in segments. On-target seeds cannot contain sequencing errors, therefore they must be completely embedded inside a segment. So the sizes of the segments indicate whether the read contains an on-target seed or not.

**Figure 6:**
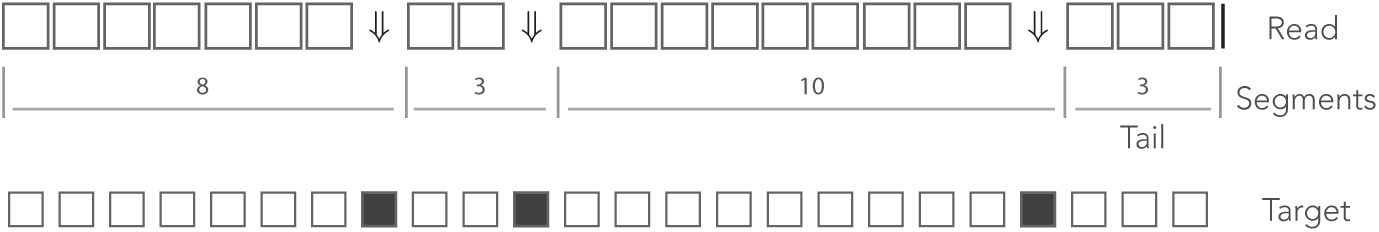
The mismatch encoding. An example read is represented in the mismatch alphabet. The symbol ⇓ represents a mismatch against the target (an erroneous nucleotide) and the symbol □ represents a match (a correct nucleotide). The symbol | is appended to the end of the read. The symbolic sequence of the target is represented below, where an open square stands for a match and a closed square stands for a mismatch.

The probability of occurrence of the symbol ⇓ is *p* (the error rate of the sequencer) and the probability of occurrence of the symbol □ is thus 1−*p* = *q*. Both symbols have size 1, so their respective weighted generating function are *pz* and *qz*. Using the rule for concatenation, we see that the weighted generating function of a segment with *i* symbols □ is (*qz*)^*i*^*pz*. The symbol has size 0, so the weighted generating function of a tail segment with *i* symbols □ is (*qz*)^*i*^.

The key insight is that reads without on-target seed are exactly the reads that are made of segments with fewer than *γ* symbols □. The weighted generating function of such segments is *pz*⋅ (1+*qz* +…+(*qz*)^*γ−*1^), and that of the tail is 1+*qz* +…+(*qz*)^*γ−*1^. This gives a construction plan that can be encoded in a transfer matrix.

There are two kinds of objects: the ⇓ segments and the tails, so the dimension of the transfer matrix is 2×2. A ⇓ segment can be followed by another ⇓ segment or by the tail. The tail cannot be followed by anything. The expression of the transfer matrix *M*_0_(*z*) is thus

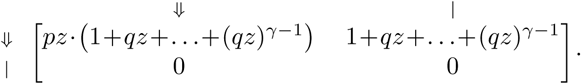

In the representation above, the different types of objects are identified by their terminator. The terminators are written in the margins of the transfer matrix for readability.

We can compute the matrix *M*_0_(*z*)+*M*_0_(*z*)^2^+…=*M*_0_(*z*)⋅(*I − M*_0_(*z*))^*−*1^ and extract the entry of interest, which is the top right term — associated with terminators ⇓ and |. To see why, observe that every read can be prepended by ⇓ segments and only by those (not by a tail). Thus, reads are precisely the sequences of segments that can follow the symbol ⇓ and that are terminated by a tail, whose weighted generating function is the top right entry of the matrix. It is easy to check that this term is equal to

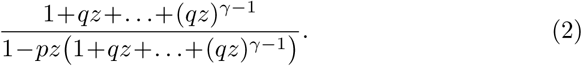

This function can be expanded as a Taylor series *a*_0_ +*a*_1_*z* +*a*_2_*z*^2^ +…, but the coefficients *a*_0_, *a*_1_, *a*_2_, … are unknown. By construction, *a*_*k*_ is the quantity of interest, *i.e.* it is the probability that a read of size *k* does not contain an ontarget seed, so we now need to extract the coefficients from expression (2). There are several methods to do so; the one we choose here is to build a recurrence equation to compute *a*_*k*_ from the previous terms of the series *a*_0_, *a*_1_, …, *a*_*k*−1_. For this, we observe that for *z*≠ 1/*q*, equation

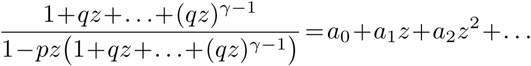

is equivalent to

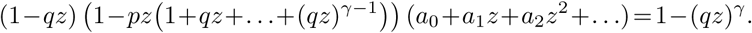

Balancing the terms of same degree on both sides of the equation leads to the following relationship between the coefficients of the series:

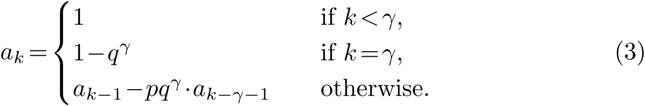

These results are consistent: for *k<γ*, the read is too short so the probability that it contains no seed is 1; for *k* = *γ*, the read contains a seed if and only if it has no error, which occurs with probability *q^γ^*.

The terms of interest can be computed recursively using expression (3). This approach is very efficient because every iteration involves at most one multiplication and one subtraction. Also, the default floating-point arithmetic on modern computers gives sufficient precision to not worry about numeric instability for the problems considered here (we rarely need to compute those probabilities for reads above 500 nucleotides).

This example shows how the symbolic approach yields a non-trivial and yet simple algorithm to compute the probability that a read of size *k* does not contain an on-target exact seed.

### 3.5 Example 2: on-target skip seeds

We go through another essential example, namely we devise a method to compute the probability that a read contains no on-target skip seed. Using the same strategy as in the previous example, we start by recoding the reads using a specialized alphabet to solve this problem.

As in the previous example, we need to know whether a nucleotide is a sequencing error, but this time we also need to know its phase in the repeated cycles of skipped positions. For this, we define the skip-*n* alphabet 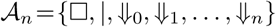. Again, the symbol □ represents a correct nucleotide and the symbol | is a terminator added at the end of the read. The symbols 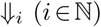 represent sequencing errors and *i* indicates the number of nucleotides till the next non-skipped position (*i.e. i* = 0 for nucleotides immediately before a non-skipped position and *i* = *n* for nucleotides at a non-skipped position).

As per definition 7, segments in this alphabet are sequences of 0 or more symbols □ followed by any of the symbols ⇓_*i*_ or by the symbol |. Given that this decomposition is unique, we can again view a read as a sequence of segments with a tail. The example of Fig. 6 is shown again in Fig. 7, where segments in the mismatch alphabet have been replaced by segments in the skip-3 alphabet. The probability of occurrence of a sequencing error is *p*, so every symbol ⇓_*i*_ has the same weighted generating function *pz* — implicitly assuming that the next non-skipped position is at distance *i*, otherwise the weighted generating function is 0. The weighted generating function of the symbol □ is again *qz*, and so the weighted generating function of a segment with *i* symbols □ is (*qz*)^*i*^*pz*. Likewise, the weighted generating function of a tail with *i* symbols □ is (*qz*)^*i*^.

**Figure 7:**
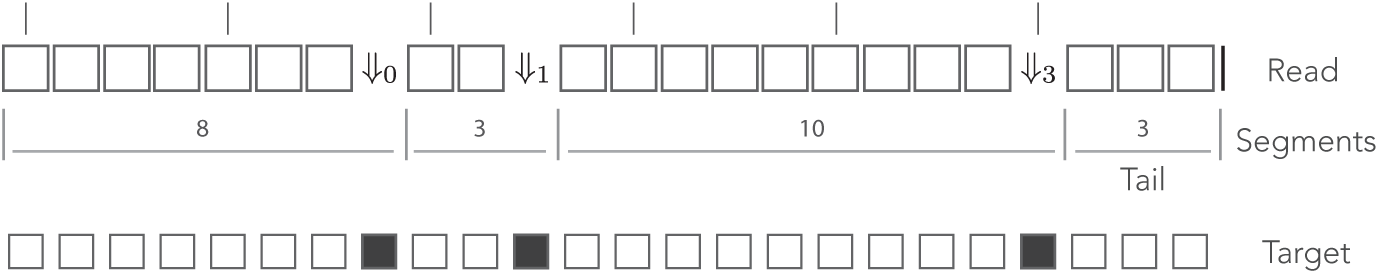
The skip encoding. The read of Fig. 6 is represented in the skip-3 alphabet. The symbols ⇓_*i*_ (*i*=1, 2, 3) represent mismatches against the target (they are erroneous nucleotides) and the symbol □ represents a match (it is a correct nucleotide). The vertical bars indicate non-skipped positions (the potential start of a seed). The number *i* in ⇓_*i*_ indicates the number of nucleotides till the next non-skipped position. For *i* = 0 the next nucleotide is not skipped. Other features are as in Fig. 6.

Reads that do not contain any on-target skip seed can contain segments with *γ* or more symbols □, so this case is more complex than the previous one. For instance, if there is a sequencing error *i* nucleotides before the next non-skipped position, the status of the next *i* nucleotides does not have any influence on the presence of an on-target seed. Indeed, there is no on-target seed upstream of the error (by construction) and the next seed is scheduled to start *i* nucleotides downstream, so there can be up to *γ* + *i*−1 symbols □ in a row. In order to enforce the absence of on-target seeds, we thus have to adjust the maximum size of the segments depending on the preceding terminator.

In the skip-*n* alphabet, reads are composed of segments terminated by the *n*+1 symbols ⇓_0_, ⇓_1_, …, ⇓_*n*_, plus the tail. The dimensions of the transfer matrix are thus (*n*+2)*×*(*n*+2).

As mentioned above, after the terminator ⇓_*i*_, we can append segments with up to *γ* + *i*−1 symbols □. The terminators of those *γ* + *i* possible segments are distributed according to the laws of the arithmetic modulo *n* + 1. When *i>* 0, for instance, if the next segment has no symbol □ then it is terminated by ⇓_*i−*1_. But the same goes if the segment has *n* + 1 symbols □ (assuming *n*+1 *< γ*). Specifying the transfer matrix thus involves a substantial amount of bookkeeping. One can check that the expression of the transfer matrix *M*_*n*_(*z*) is

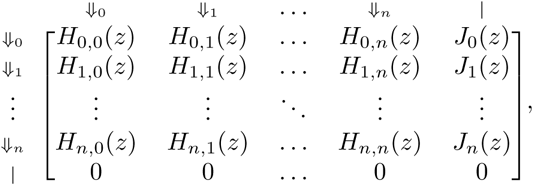

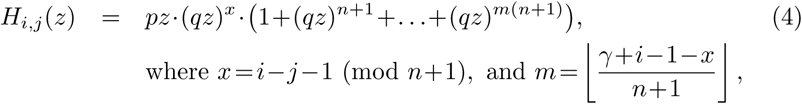

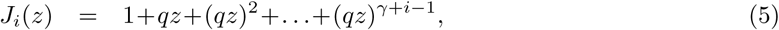

where └…┘ denotes the “floor” function.

The weighted generating function of interest is the top right entry of the matrix *M*_*n*_(*z*)+*M*_*n*_(*z*)^2^ +…= *M*_*n*_(*z*)⋅(*I − M*_*n*_(*z*))^*−*1^. To see why, observe that, at the start of every read, the next nucleotide is a non-skipped position, so every read can be prepended by ⇓_0_ segments and only by those. Thus, reads are precisely the sequences of segments that can follow the symbol ⇓_0_ and that are terminated by a tail, whose weighted generating function is the entry of the matrix associated with terminators ⇓_0_ and |.

*M*_*n*_(*z*) is too complex to allow a closed expression of *M*_*n*_(*z*)⋅(*I − M_n_*(*z*))^*−*1^ to be computed. We return to the definition *M*_*n*_(*z*)+*M*_*n*_(*z*)^2^ +…and observe that the terms *M*_*n*_(*z*)^*k*+2^, *M*_*n*_(*z*)^*k*+3^, … have no influence on the coefficients of the weighted generating function *a*_0_, *a*_1_, …, *a*_*k*_.

Indeed, a read of size *k* has at most *k*+1 segments (including the obligatory tail). Since *M* (*z*)^*s*+1^ contains the weighted generating functions of reads with exactly *s*+1 segments including the tail, *a*_*k*_ cannot depend on *M* (*z*)^*k*+2^*, M* (*z*)^*k*+3^, …. More formally, we can prove by induction that all the entries of *M* (*z*)^*k*^ are divisible by *z*^*k*−1^, showing that the contribution of *M* (*z*)^*k*+2^ + *M* (*z*)^*k*+3^ + … to *a*_0_ +*a*_1_*z*+ +*a*_*k*_*z*^*k*^ is strictly 0.

So we can compute the matrix *M*_*n*_(*z*)+ *M*_*n*_(*z*)^2^ + …+ *M*_*n*_(*z*)^*k*+1^, extract the entry of interest and then compute the terms of the Taylor expansion up to order *k*. But we can do better: since we are only interested in the coefficients up to order *k*, we can perform all algebraic operations on truncated polynomials of order *k*, *i.e.* we discard the coefficients of order *k*+1 or greater when multiplying two polynomials.

But we can do even better: a read with *s*+1 segments contains *s* errors, so all the entries of *M*_*n*_(*z*)^*s*+1^ are dominated by *p*^*s*^ and they rapidly converge to 0 as *s* increases. Instead of computing the matrix *M*_*n*_(*z*)+*M*_*n*_(*z*)^2^+…+*M*_*n*_(*z*)^*k*+1^, we can interrupt the summation after a certain power of *M*_*n*_(*z*) because the terms become negligible.

The number of errors *X* in a read of size *k* has a Binomial distribution *X ~ B*(*k, p*). From [34] we can bound the probabilities of the tail with the expression

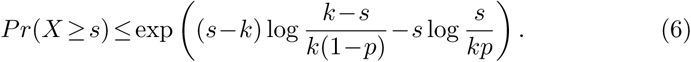

Using the formula above, we can thus bound the probability that a read has *s* + 1 segments or more. We compute *M*_*n*_(*z*)+ *M*_*n*_(*z*)^2^ + …+ *M*_*n*_(*z*)^*s*+1^ where the weighted generating functions have been replaced by truncated polynomials and we extract the top right entry. When the bound is lower than a set fraction *ε* of the current value of *a*_*k*_, we stop the computations. Typically *ε* = 0.01 so this method ensures that the probabilities that a read of size *k* has no on-target skip seed are accurate to within 1%.

***Remark 1.** Observe that when n*=0 *the matrix M*_*n*_(*z*) *is identical to the matrix M*_0_(*z*) *of section 3.4. This is consistent with the fact that exact seeds are skip-*0 *seeds*.

## 4 Off-target exact seeds

We now turn our attention to the problem of computing the probability that the seeding process is off target when using exact seeds — recall from section 1.3 that off-target seeding means that the candidate set contains a duplicate but not the target.

If there is *N* = 0 duplicate, seeding cannot be off target, it can only be on target or null. So from here we assume that the target has *N ≥*1 duplicates. Let *S*_0_ denote the event that there is an on-target seed and let *S*_*j*_ denote the event that there is a seed for the *j*-th duplicate. We are thus interested in computing 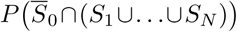, where 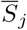 denotes the complement of the event *S*_*j*_. For this, we first observe that

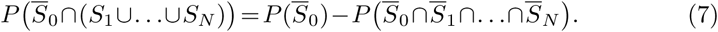

Since the duplicates are assumed to evolve independently of each other and through the same mutagenesis process, the events are conditionally independent (we do not have full independence because *S*_0_ only depends on sequencing errors whereas the events *S*_*j*_ depend on sequencing errors and on the divergence between duplicates). We can thus write

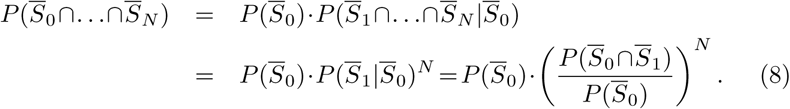

Combinging the two equations above, we obtain

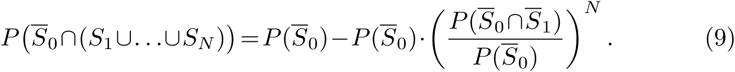

Hence, the probability that seeding is off target is a function of two quantitites. We have already seen how to compute 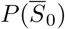 in section 3.4 using recursive expression (3). We now need to find a way to compute 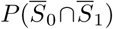.

### 4.1 The dual encoding

The event 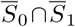 is that the read contains no seed for either the target or for the first duplicate — numbering is arbitrary here, we simply chose a duplicate and stick to it. As illustrated in section 3, we first recode the reads using a specialized alphabet to simplify the problem.

It will be useful to consider a more general problem where we have two sequences of interest labelled + and *−*. The + sequence stands for the target and that the *−* sequence stands for its duplicate. We then define the dual alphabet 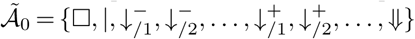. The symbols 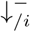 signify that the nucleotide is a mismatch against the *−* sequence only, the symbols 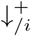 signify that it is a mismatch against the + only, and the symbol ⇓signifies that it is a mismatch against both. As before, every other nucleotide is replaced by the symbol □, and the terminator | is appended to the end of the read. We again define reads as sequences of segments, except that now the terminators are the symbols 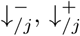 and ⇓. The tail, as usual, is terminated by the symbol |.

The number *i* in the symbol 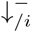 indicates the match length of the + sequence. Likewise, the number *i* in the symbol 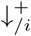 indicates the match length of the *−* sequence. For instance, the symbol 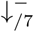 indicates that the nucleotide is a mismatch against the − sequence, that it is a match for the + sequence, and that the six previous nucleotides were also a match for the + sequence. The terminators thus encode the local state of the read.

Fig. 8 shows an example of read in the dual encoding. The + and − sequences are represented below the read, with matches represented as open squares and mismatches as closed squares. It is visible from this example that symbols 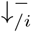 and 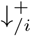 alternate whenever the mismatches hit different sequences. The symbol ⇓ occurs only when a nucleotide is a double mismatch.

**Figure 8:**
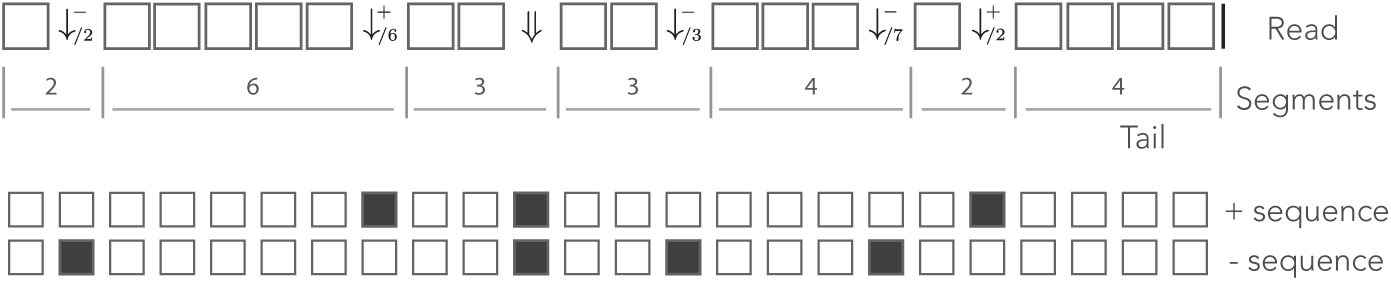
Example of dual encoding. An example of read is represented in the dual alphabet. The symbols 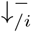 represent a mismatch against the *−* sequence, the symbols 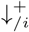 represent a mismatch against the + sequence, and the symbol ⇓ represent a mismatch against both. The index *i* is the match length of the sequence that is not mismatched. The symbolic + and *−* sequences are represented below, where an open square stands for a match and a closed square stands for a mismatch.

We assume that for each nucleotide, *a* is the probability that the read matches both sequences, *b* is the probability that it matches only the + sequence, *c* is the probability that it matches only the − sequence and *d* is the probability that it matches none. Obviously *a*+*b*+*c*+*d* = 1. These definitions allow us to specify the weighted generating function of the symbols and of the segments. To specify the transfer matrix, we only have to make sure that we eliminate the combinations that create a match of size *γ* or more for any of the two sequences. For notational convenience, we define

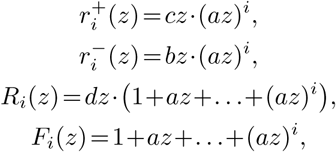

and we can verify that the expression of the transfer matrix 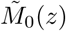 is

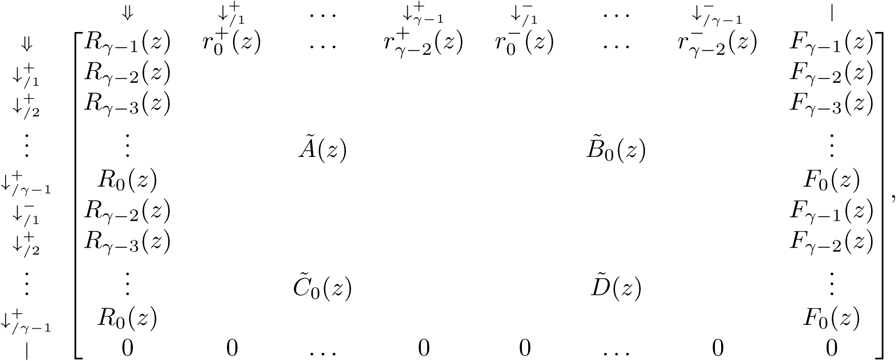

where 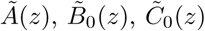 and 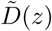 are matrices of dimensions (*γ* − 1)×(*γ* ℒ 1) that are defined as

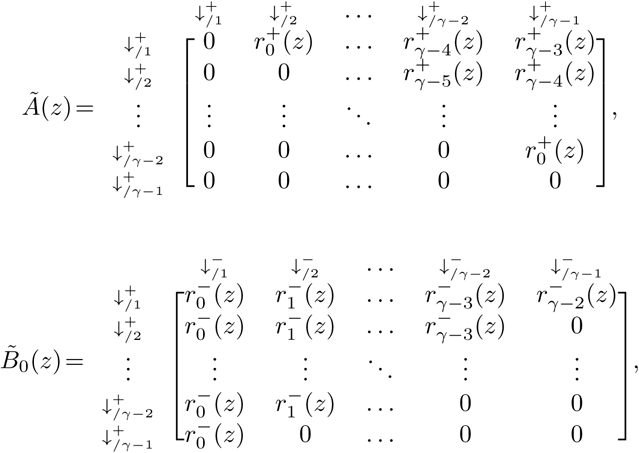

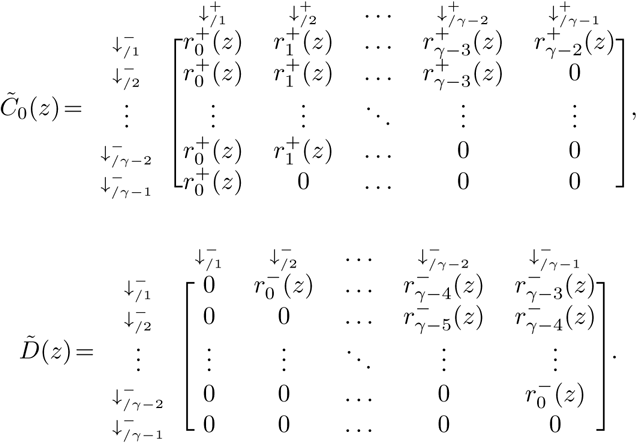

The term of interest is the top right entry of 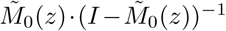. To see why, observe that every read can be prepended by ⇓ segments and only by those (every other terminator would imply that one of the two sequences has a nonzero match size at the start of the read). Thus reads are precisely the sequences of segments that can follow the symbol ⇓ and that are terminated by a tail, whose weighted generating function is the top right entry of the matrix.

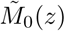 is too complex to allow a closed expression of 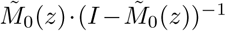 to be computed. It is actually easier to proceed as in section 3.5 and to compute the powers of 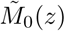 (using the arithmetic of truncated polynomials) up to a given threshold value. Since each segment except the tail contains a mismatch against at least one sequence, we define 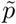 as the upper bound on the probability of a mismatch 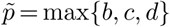. The updated formula (6) now gives an upper bound of the probability that a read of size *k* contains *s* or more mismatches as

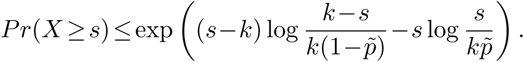

Now returning to the problem of computing 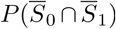, the + sequence is interpreted as the target and the − sequence as the duplicate. Based on the assumptions of the error model presented in section 3.1, this implies that *a* =(1 − *p*)(1 − *µ*), *b* =(1 − *p*)*µ*, *c* = *pµ*/3, and *d* = *p*(1 − *µ*/3).

With these values, we can fully specify the matrix 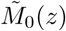, and compute its powers until the upper bound is lower than a set fraction *ε* of the current value of *a*_*k*_, which gives an accurate estimate of 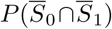.

***Remark 2.** Observe that expression (8) is the probability of null seeding. The method described above thus allows us to compute this probability at no additional cost. This probablity is less relevant than the probabilities that the seeding process is on target or off target, but at times, it may be useful to know the probability that a read is not mappable, especially when reads are relatively short*.

### 4.2 Illustration

We illustrate the strategy delineated above for reads of size *k* = 50 sequenced with an instrument with error rate *p*=0.01, when using exact seeds of size *γ* =17.

Fig. 9 shows the result for a number of duplicates *N* from 1 to 10 and for a divergence rate *µ* from 0 to 0.20. The first observation is that the probability that seeding is off target increases with *N*. This is clearly visible from expression (9). This can also be understood intuitively because the probability of null seeding decreases when there are more potential candidates, and since the probability of on-target seeding does not change, the probability of off-target seeding must increase.

**Figure 9:**
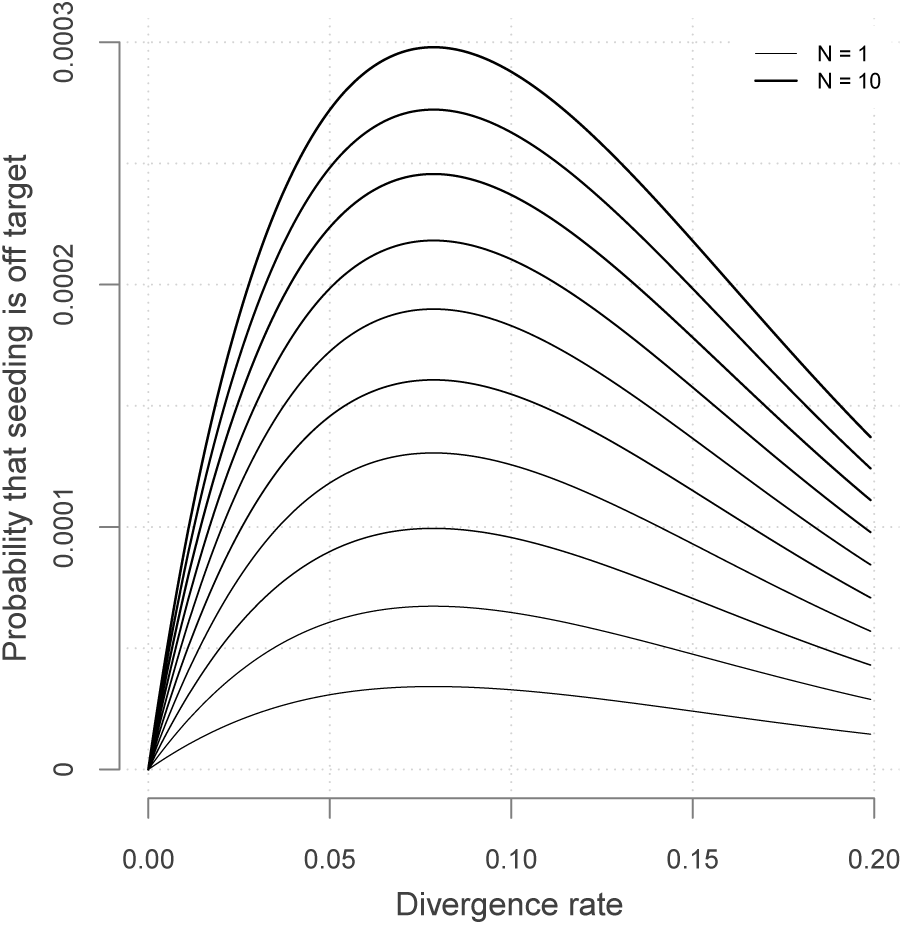
Off target seeding probabilities (exact seeds). The curves show the probability that seeding is off target for exact seeds of size *γ* =17 in reads of size *k* =50 nucleotides sequenced with an error rate *p* = 0.01. Each line shows a different number of duplicate sequences *N* from 1 to 10 and the x-axis shows the divergence rate *µ*, defined as the probability that a given duplicate differs from the target at any given position.

The second observation is that there exists a “worst” value of *µ* situated around 0.08. When *µ* is much smaller, the sequences of the duplicates are close to that of the target so it is unlikely that the candidate set contains one but not the other. When *µ* is much larger, the sequences of the duplicates are far from that of the target and they are unlikely to be in the candidate set. In expression (9), the only term that depends on *µ* is 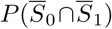, and it is obvious that the minimum of expression (9) corresponds to the maximum of 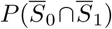. This is why the worst value of *µ* is the same for every *N*.

For comparison purposes, it is important to notice the order of magnitude of the probabilities. Up to *N* = 3 duplicates, off-target seeding has probability lower than 10^*−*4^ and for *N* =10 duplicates the upper bound is 3⋅10^*−*4^. This sets a baseline to compare with more elaborate seeding methods.

## 5 Off-target skip seeds

To compute the probability that the seeding process is off target when using skip seeds, we observe that the logic of section 4 can be transposed with few modifications. In particular, the probability can be computed through expression (9), where *S*_0_ is the event that the read has a *skip* seed for the target and *S*_1_ that it has a *skip* seed for the first duplicate.

We have already seen how to compute 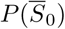 in section 3.5, we now need to find a way to compute 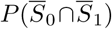 when using skip seeds.

### 5.1 The skip dual encoding

Once again, we start by recoding the reads in a specialized alphabet. We define the skip-*n* dual alphabet 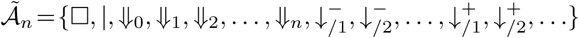. The symbols □, *|*, 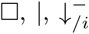 and 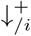 have the same meaning as in the dual alphabet of section 4.1. The symbols ⇓_*i*_ indicate that both sequences have match length 0 and that the next non-skipped position is *i* nucleotides downstream.

Fig. 10 shows the read from Fig. 8 represented in the skip-3 dual encoding. It is important to note several differences with Fig. 8. The first is that the symbols ⇓_*i*_ are not always associated with double mismatches. For instance, the symbol ⇓_0_ on the right side of the read corresponds to a mismatch for the + sequence only. This happens whenever the + and the − sequences are mismatched in the same interval between non-skipped positions (the mismatches do not need to be on the same nucleotide).

**Figure 10:**
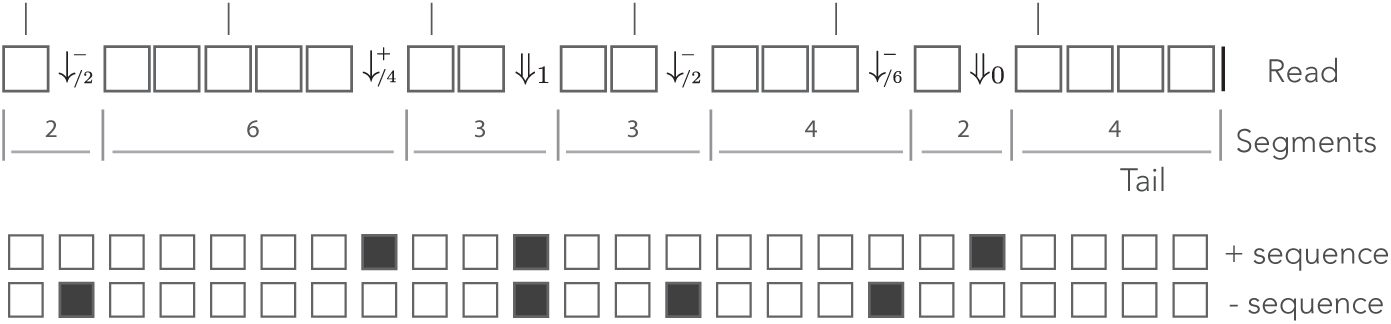
Example skip dual encoding. The read of Fig. 8 is represented in the skip dual alphabet. The vertical bars above the read indicate non-skipped positions. The symbols 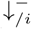 and 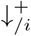 have the same meaning as in the dual alphabet. The symbols ⇓_*i*_ indicate that both sequences have match length 0 and that the next non-skipped position is located *i* nucleotides downstream. Other features are as in Fig. 8.

As in the case of the skip encoding, a fair amount of bookkeeping is required to specify the weighted generating functions of interest. We reuse the quantities *a*+*b*+*c*+*d* =1 and give them the same meaning as in section 4.1. For notational convenience, we define

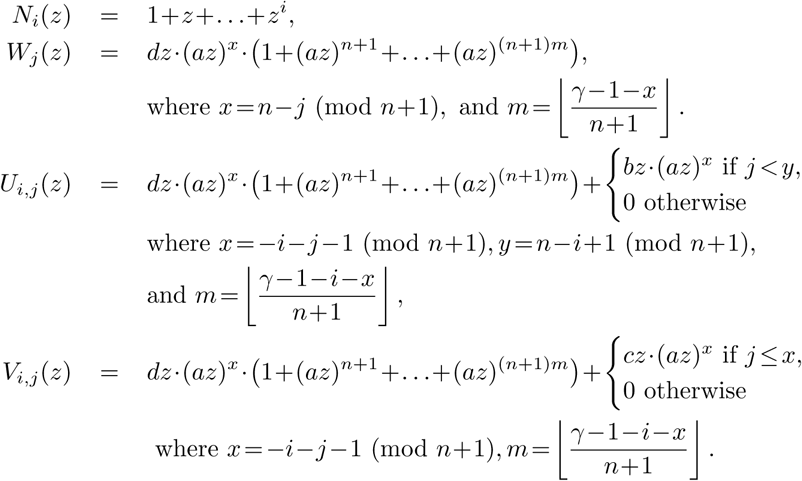

With these definitions, one can check that the expression of the transfer matrix 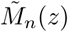 is

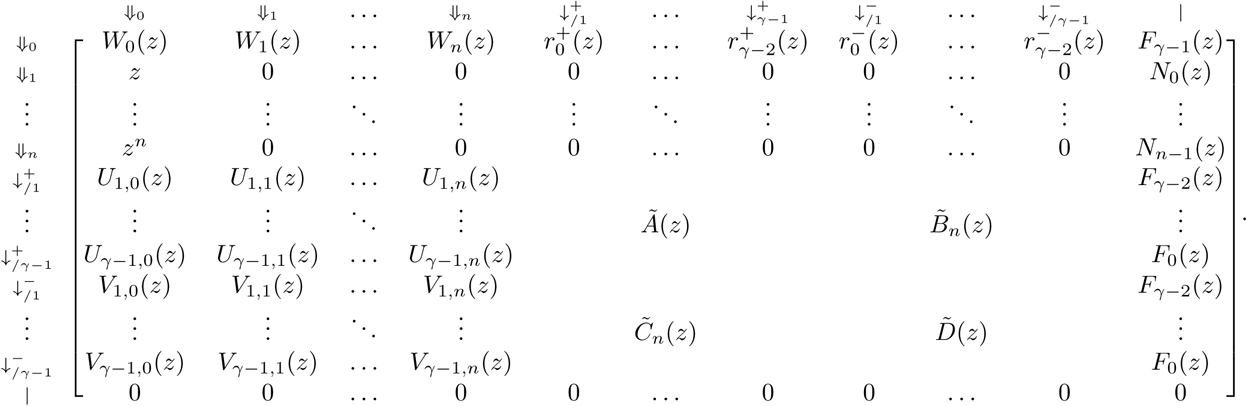

The matrices 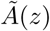 and 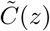 in the expression of 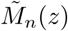 are the same as in section 4.1. The matrices 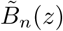 and 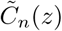 are defined as

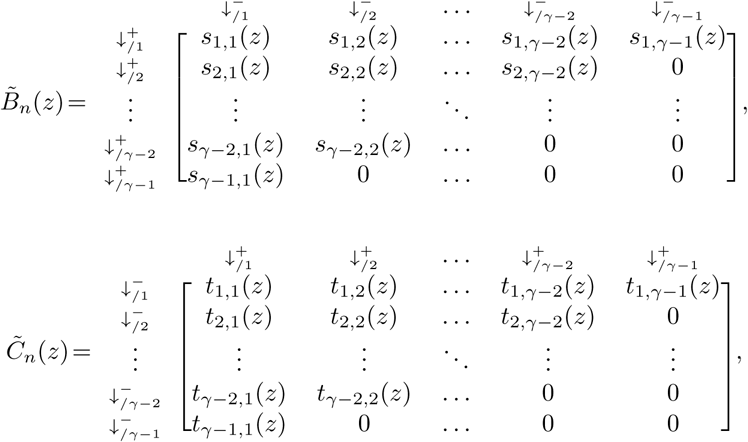

with

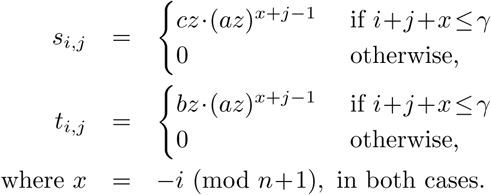

The computation is performed exactly as described in section 4.1. We compute the successive powers of 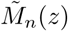 in the arithmetic of truncated polynomials and stop the iterations using the same criterion. So there is no change other than the definition of the transfer matrix.

***Remark 3.** Observe that when n* = 0 *the matrix* 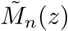 *is identical to the matrix* 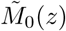 *of section 4.1, again consistent with the fact that exact seeds are skip-*0 *seeds. The same applies to* 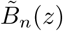 *and* 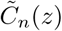.

### 5.2 Illustration

We illustrate the strategy delineated above Using the same settings as in section 4.2 (*k* = 50, *p* = 0.01 and *γ* = 17), except that we use skip-9 seeds instead of exact seeds.

Fig. 11 shows the result for a number of duplicates *N* from 1 to 10 and for a divergence rate *µ* from 0 to 0.20. The curves have the same general aspect as those of Fig. 9. The probability that seeding is off target increases with *N*.

**Figure 11:**
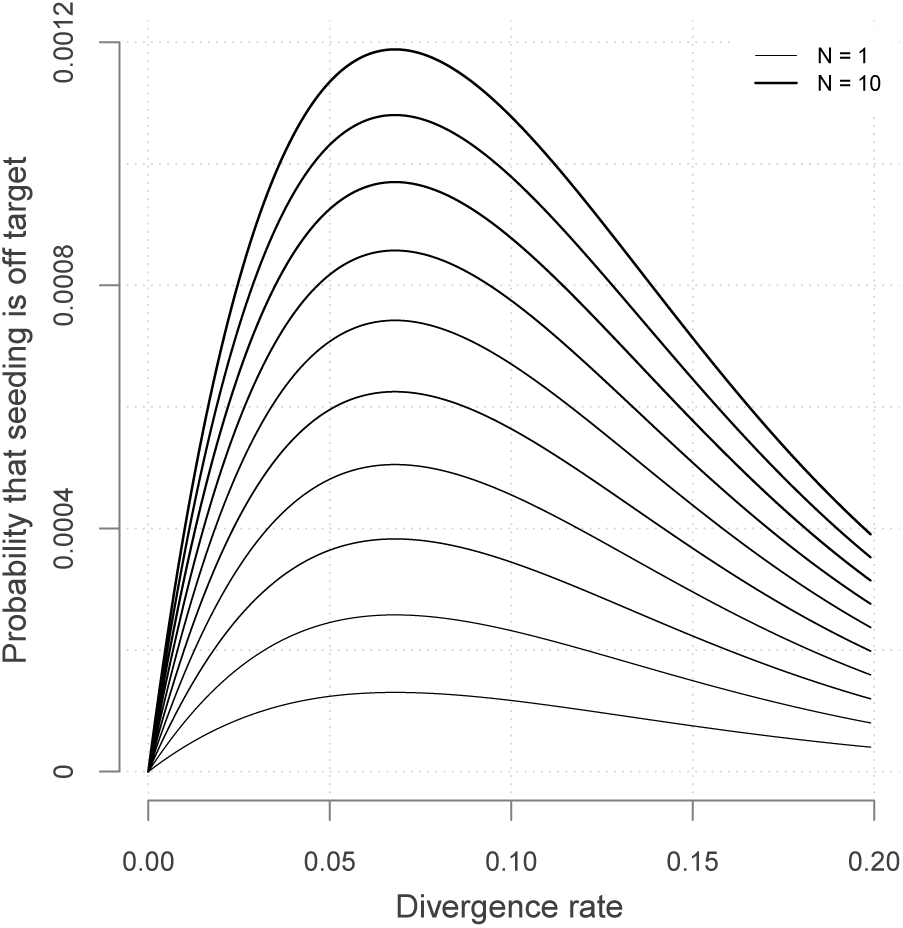
Off target seeding probabilities (skip seeds). The curves show the probability that seeding is off target for skip-9 seeds of size *γ* =17. Read size and error rate are the same as in Fig. 9, *i.e. k* = 50 and *p* = 0.01. Each line shows a different number of duplicate sequences *N* from 1 to 10 and the x-axis shows the divergence rate *µ*, defined as the probability that a given duplicate differs from the target at any given position.

There is again a worst value of *µ*, because the maximum of 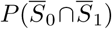 minimizes expression (9) for every value of *N*. But the value is different from that of Fig. 9 because 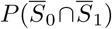 is computed for skip-9 seeds instead of exact seeds (here the worst value of *µ* is approximately 0.07).

Importantly, Fig. 11 reveals that skipping 9 nucleotides increases the chances that seeding is off target by a factor approximately 4 (compare with Fig. 9). It is not obvious why this should be the case: skip seeds reduce the probability of on-target seeding (skipping positions implies fewer on-target seeds) but increase the probability of null seeding (skipping positions implies fewer seeds overall). So the net effect on off-target seeding is not intuitive.

And indeed, there are cases that skip seeds are *less* prone to off-target seeding than exact seeds. This is the case for instance when *p* = 0.1, where the ranking is inverted for the whole range of values considered in Fig. 11.

This information is critical for choosing the best seeding strategy. The offtarget seeding probability is not the only criterion, though. Equally important considerations are the probability of on-target seeding and the computational resources required to implement a particular seeding strategy. The benefit of a theory to compute seeding probabilities is to have access to this knowledge.

## 6 Off-target MEM seeds

MEM seeds are substantially more complex than exact seeds and skip seeds because we need to take into account all the duplicates in the combinatorial construction.

### 6.1 Hard and soft masking

We first introduce two important notions that will be the key of understanding the behavior of MEM seeds.

#### Definition 8.

*At a given position of the read, a duplicate is a* hard mask *if its match length on the left side is strictly longer than the match length of the target. A duplicate is a* soft mask *if it has the same match length as the target*.

Fig. 12 gives a graphical intuition of hard and soft masks. It is important to bear in mind that hard and soft masks depend on the position of interest: a sequence can be a mask at the left end of the read and not at the right end, or the opposite.

Hard and soft masks explain the counter-intuitive properties of MEM seeds. For instance, in Fig. 4 the target cannot be discovered because every nucleotide of the read has a hard mask. In Fig. 5, the target could be discovered if the read were shorter because a hard mask would turn into a soft one.

From the definition, we see that the last nucleotide of every strict on-target MEM seed is always unmasked. Conversely, an unmasked nucleotide always belongs to exactly one strict on-target MEM (not necessarily a seed because the size of the MEM can be less than *γ*). Also, the last nucleotide of every shared on-target MEM seed is always soft-masked, but a soft-masked nucleotide does not always belong to a shared on-target MEM.

Since hard and soft masks inform us about the positions of on-target MEM seeds, we construct an alphabet that encodes the masking status of the nucleotides.

**Figure 12:**
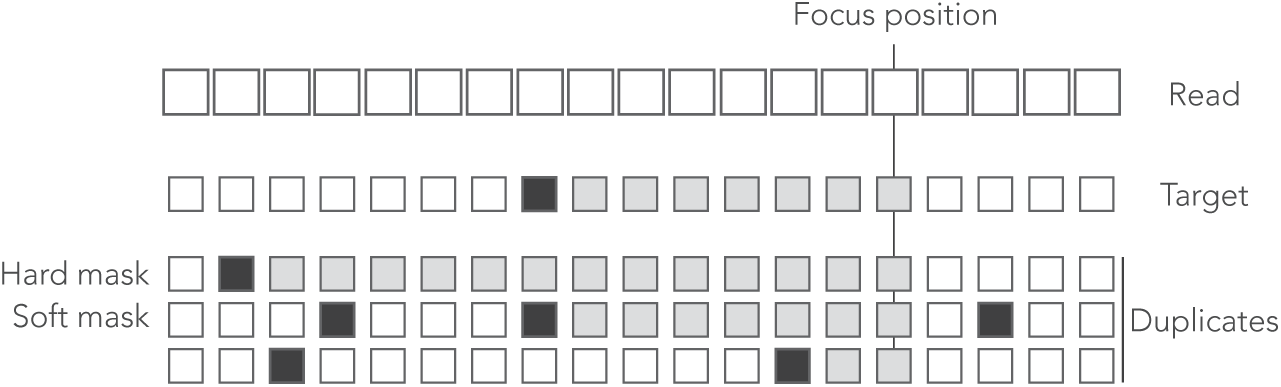
Example of hard and soft masks. Genomic sequences are shown below a read where open squares represent nucleotides. In the sequences, the open squares represent matches and the closed squares represent mismatches. The nucleotides contributing to the match length are represented as grey boxes. At the focus position, the match length of the target is 7. The first duplicate is a hard mask because its match length is 13 *>* 7. The second duplicate is a soft mask because its match length is 7, as the target. The third duplicate is not a mask because its match length is 2 *<* 7.

### 6.2 The MEM alphabet

As before, we recode the reads as sequences of letters from a specialized alphabet called the MEM alphabet 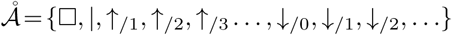.

The symbols *↓*_*/m*_ indicate that the nucleotide is a sequencing error and *m≥*0 is the number of duplicates that *match* the nucleotide. Since a sequencing error is always a mismatch against the target, the symbol *↓*_/0_ indicates that the nucleotide is a mismatch against *every* sequence. The symbols ↑_*/i*_ indicate a change in masking status: the nucleotide is not masked but the previous is — this happens when all the duplicates fail to extend beyond this position. The index *i* ≥ 1 is the number of nucleotides since the last mismatch or since the beginning of the read. All the other nucleotides are represented by the symbol □ and the symbol *|* is appended to the end of the read as usual.

Note that in the symbols *↓*_/*m*_ and *↑*_/*i*_, the numbers *m* and *i* have different meanings. In the symbol *↓*_/*m*_, the index *m* is a number of sequences (0*≤m≤N* where *N* is the number of duplicates); in the symbol ↑_*/i*_, the index *i* is a number of nucleotides. Fig. 13 shows the encoding of a read in the MEM alphabet.

**Figure 13:**
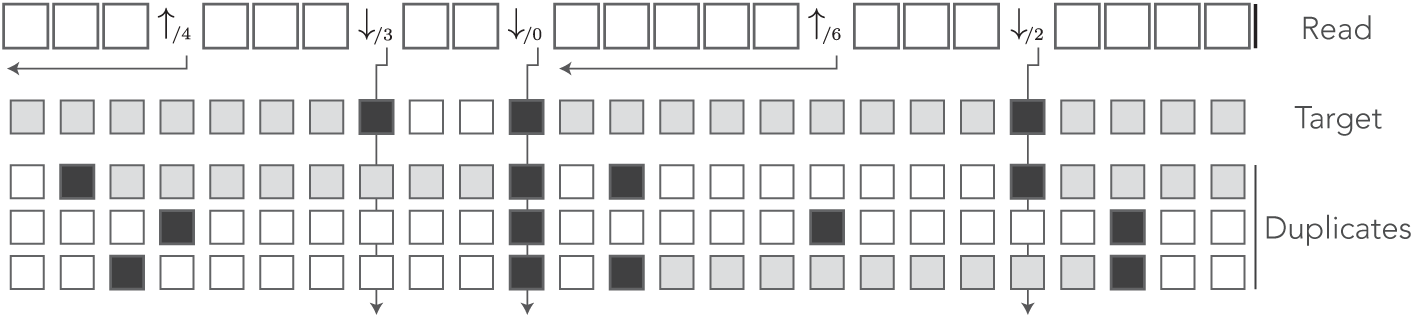
The MEM encoding. The read of Fig. 3 is represented in the MEM alphabet. The arrows departing from the numbers help understand their meaning. The symbol *↓*_/*m*_ is indexed by the number *m* of locations that match the nucleotide. The symbol *↑*_/*i*_ is indexed by the number *i* of nucleotides from the last error or from the beginning of the read. The grey squares in the symbolic sequences represent MEM seed matches. Other features are as in Fig. 8.

The MEM alphabet captures the masking status of the nucleotide: the symbol *↓*_/*m*_ indicates that the nucleotide has *m* hard masks and *N −m* soft masks. The symbols ↑_*/i*_ indicate that the nucleotide is unmasked and that the previous nucleotide is masked.

In the MEM alphabet, strict on-target MEM seeds are the longest stretches of symbols containing some symbol ↑_*/i*_ and not containing any symbol ↓_*/m*_. Indeed, such a stretch is a match for the target because it does not contain any symbol *↓*_/*m*_, it only matches the target because it contains at least one unmasked nucleotide (marked by *↑*_/*i*_), and it cannot be extended because it is flanked by sequencing errors (symbols *↓*_/*m*_) or by the ends of the reads. Note that there is exactly one symbol *↑*_/*i*_ per strict on-target MEM seed, and therefore two symbols ↑_*/i*_ must be separated by at least one symbol ↓_*/m*_.

Shared on-target MEM seeds are the longest stretches of symbols □ flanked by *↓*_/0_, or by the ends of the read. Indeed, such a stretch is a MEM seed because it matches the target and it cannot be extended (↓_*/*0_ is a mismatch against every sequence). Also, it cannot be a strict on-target MEM seed because it does not contain any ↑_*/i*_ symbol, so it must be a shared on-target MEM seed.

As before, the read is converted from a sequence of symbols to a sequence of segments that consist of 0 or more symbols □ followed by a terminator. We then specify the weighted generating functions of those segments and fill the transfer matrix 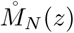 of the reads that do not contain an on-target MEM seed. We introduce the terms of the matrix by increasing order of complexity.

### 6.3 Segments following ↑_/*i*_

A segment terminated by ↑_*/i*_ is the beginning of a strict on-target MEM of size at least *i*. The MEM reaches the next sequencing error or the end of the read, so the number of symbols □ in the next segment must be at most *γ−i−*1 and it must be terminated by a ↓_*/m*_ symbol or by the tail terminator |.

The following definition will simplify the notations.

#### Definition 9.

*The probability that a symbol is* ↓_/*m*_ *given that the nucleotide is a read error is*

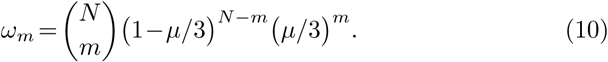

The expression for *ω_m_* is exact: if the nucleotide is an error, the symbol is ↓_*/m*_ for some *m* between 0 and *N*. Each duplicate is a match with probability *µ/*3, so *m* has a Binomial distribution with parameters (*N, µ/*3).

On this segment, the matches between the read and the duplicates are irrelevant, so the weighted generating function of a symbol □ is simply *qz* (recall that *p* =1*−q* is the probability of a sequencing error). The weighted generating function of the terminator *↓*_/*m*_ is *ω*_*m*_*pz*, so the weighted generating function of the *↓*_/*m*_ segments following *↑*_/*i*_ is

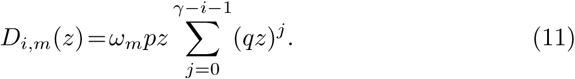

And the weighted generating function of the tail segments following *↑*_/*i*_ is

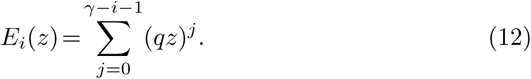

### 6.4 Segments following *↓*_/*m*_

The symbol ↓_*/m*_ signifies that the nucleotide has *m* hard masks and *N − m* soft masks. If all the masks vanish before the first read error, the next terminator will be a symbol ↑_*/j*_, otherwise it will be the symbol | or a symbol ↓_*/m*_. We separate the cases based on the terminator of the segment.

#### Case 1: the terminator is ↑_/*j*_

##### Definition 10.

*The probability that a given duplicate contains a mismatch in a sequence of j error-free nucleotides is*

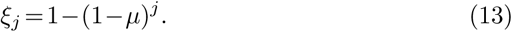

*This is the probability that a hard or soft mask vanishes within j correct nucleotides*.

The expression for *ξ*_*j*_ is exact: every nucleotide of the duplicate differs from the target with probability *µ*. In the absence of sequencing errors, this is also the probability that a nucleotide of the duplicate differs from the read. Given that there is no error, the probability that *j* nucleotides in a row are identical to the read is thus (1 − *µ*)^*j*^ and the probability that at least one of them is different is the complement 1 − (1 − *µ*)^*j*^.

With this notation, the probability that at least one of *N* masks survives a sequence of *j* error-free nucleotides is thus 1 *−* (*ξ*_*j*_)^*N*^, and the probability that there remains a mask at the *j−*1-th but not at the *j*-th error-free nucleotide is (*ξ*_*j*_)^*N*^ −(*ξ*_*j*−1_)^*N*^. From this we conclude that the weighted generating function of the segments terminated by *↑*_/*j*_ following a segment terminated by *↓*_/*m*_ is

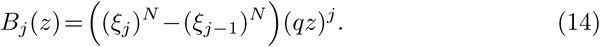

The fact that the reads have no on-target seeds imposes *j* < *γ*. Also note that this expression is the same for all symbols *↓_/m_* (it does not depend on *m*).

#### Case 2a: the terminator | comes before the ***γ***-th nucleotide

In this case there can be no on-target seed because the read finishes too early. However, we must enforce the condition that at least one of the *N* masks survives until the end, otherwise the segment would be terminated by one of the symbols *↑*_/*j*_. The weighted generating function is

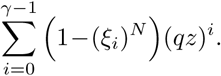

#### Case 2b: the terminator | comes after the ***γ***-th nucleotide

In this case, the soft masks do not hide the target. Even if a duplicate survives until the end of the read, there will be an on-target seed (shared in this case). To exclude on-target seeds, we must enforce the condition that at least one hard mask survives until the end of the segment (which is impossible if *m* = 0). The weighted generating function is

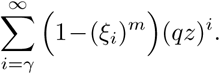

Summing the expressions from cases 2a and 2b, we find that the weighted generating function of the tail following *↓*_*/m*_ is

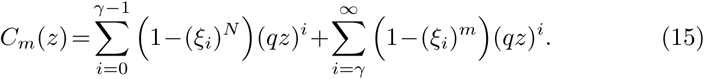

#### Case 3a: the terminator *↓*_/*n*_ comes before the *γ*-th nucleotide

In this case, there can be no on-target seed and we must only exclude the terminators ↑_*/j*_. As we have seen above, this implies that at least one of the *N* masks survives until the terminator. For a read of size *j* +1, this occurs with probability 1 − (*ξ*_*j*_)^*N*^. Including the terminator and summing over *j* +1≤*γ*, we see that the weighted generating function is>

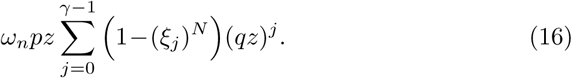

#### Case 3b: the terminator *↓*_/*n*_ comes after the *γ*-th nucleotide

This case is by far the most convoluted. Since the segment contains at least *γ* error-free nucleotides, we must enforce the condition that it does not contain an on-target seed. This will be the case if any of the two following conditions is validated: *i*) at least one hard mask covers all the error-free nucleotides, or *ii*) all the hard masks vanish but at least one soft mask covers the whole segment (including the terminator).

The two conditions are mutually exclusive by construction. They are graphically represented in the diagram below. The left panel corresponds to case *i*) and the right panel to case *ii*). The top row represents the target, and the bottom rows represent duplicates (using the same symbols as in Fig. 13).

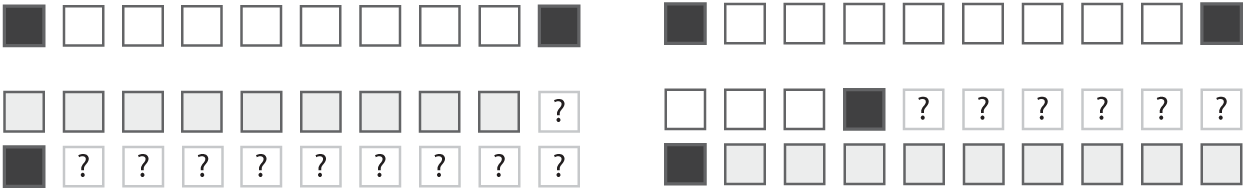

Whenever a hard mask (here the first duplicate) covers the nucleotides as shown by the grey squres in the left panel, there can be no on-target seed. The positions marked with a question mark are irrelevant, they cannot change the fact that there is no on-target MEM seed. If the hard masks vanish, as in the right panel, then we need to look at the soft masks. If a soft mask covers the whole segment as indicated by the grey squares, then there can be no on-target seed. In all other cases there is an on-target MEM seed.

For a segment of size *j*+1, condition *i*) has probability 1 −(*ξ*_*j*_)^*m*^. Summing over *j* +1 *>γ* and including the terminator, we see that the associated weighted generating function is

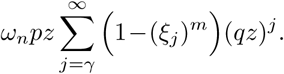

Condition *ii*) is more convoluted, so we introduce some further notations to solve this sub-case.

##### Definition 11.

*The probability that a given duplicate sequence contains a mismatch in a sequence of j error-free nucleotides followed by an error is*

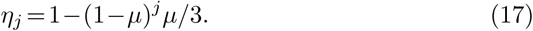

*This is the probability that a hard or soft mask vanishes within j correct nucleotides followed by a sequencing error*.

The expression for *η*_*j*_ is exact: the probability that there are *j* matches between the duplicate and the target is (1 − *µ*)^*j*^. If there are no sequencing errors, this is also the probability that there are *j* matches between the duplicate and the read. The probability that the duplicate matches the subsequent error is *µ*/3, so the probability that there are *j* +1 matches including the sequencing error is (1 − *µ*)^*j*^*µ*/3. Finally, the probability that there is a mismatch is the complement 1 − (1 − *µ*)^*j*^*µ*/3.

Let us for now consider a segment of fixed size *j* +1. From expressions (13) and (17), the probability of condition *ii*) is

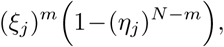

but we need to break up this term among all the possible terminators ↓_*/n*_ (0 ≤ *n* ≤ *N*) in order to fill the different entries of the transfer matrix. For this, we split this term in the number of soft masks that run until and including the terminator. From expression (17), the probability that there are *r* ≥ 1 such soft masks is

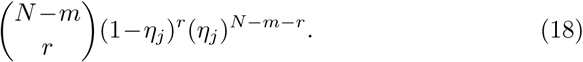

For now we consider *r* fixed; we will compute the marginal probability at the final stage. By construction, each of those *r* soft masks matches the terminator, so the total number of matches is *r* plus the number of sequences that also match the terminator, among the remaining *N − m − r* soft masks and the *m* hard masks.

Let us start with the *m* hard masks. The probability that each of them matches the terminator is simply *µ*/3.

The case of the *N − m − r* soft masks is more complicated because they can vanish precisely on the terminator — recall that in case *ii*) all the hard masks are assumed to vanish before. If the soft mask failed within the first *j* nucleotides, then the *j* +1-th nucleotide can be anything and it will match the terminator with probability *µ*/3. But if the soft mask survived the first *j* nucleotides, then it *must* fail on the *j* + 1-th and it cannot match the terminator. From expressions (13) and (17), the probability that a given soft mask fails within the first *j* nucleotides is *ξ*_*j*_/*η*_*j*_ — this is the conditional probability that it fails within the first *j* nucleotides given that it fails within the segment. Finally the probability that such a soft mask matches the terminator is *µ*/3⋅*ξ*_*j*_/*η*_*j*_.

Summing the contributions of the hard masks (*m* in total, each matching the terminator with probability *µ/*3) and of the soft masks (*N −m−r* in total, each matching the terminator with probability *µ*/3⋅*ξ*_*j*_/*η*_*j*_), the probability that the total number of matches is *n−r* appears as the convolution product

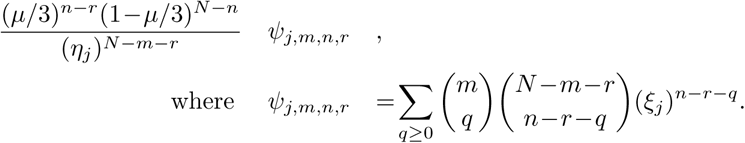

Finally, we need to compute the marginal probability over the number *r* of soft masks that survive until the end of the read. Multiplying by the probability of *r* from expression (18) and summing over *r* ≥ 1, the probability that *n* duplicate sequences match the terminator appears as

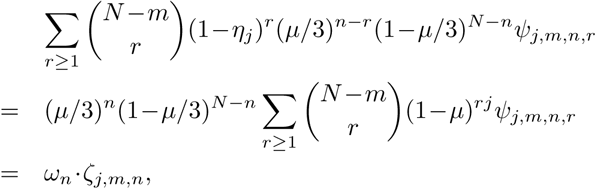

where

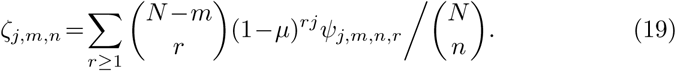

This is the probability that the terminator is the symbol ↓_*/n*_ given that the segment has size *j* + 1 > *γ*, that the first sequencing error occurs on the last nucleotide, that the preceding terminator was ↓_*/m*_ and that the *m* hard masks fail before the end of the segment.

Summing the terms from case 3a and from case 3b, we find that the weighted generating function of the *↓*_/*n*_ segments following *↓*_/*m*_ is

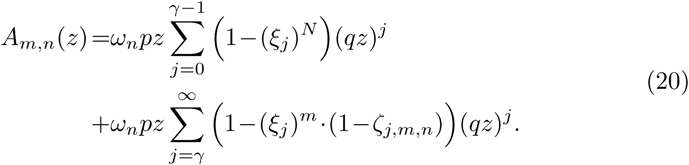

### 6.5 Expression of 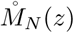

Collecting and arranging the results above, we can verify that the final expression of the transfer matrix 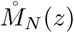 is

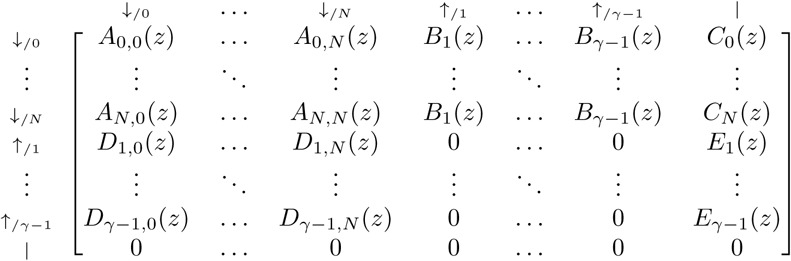

where

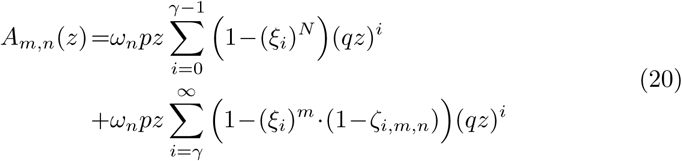

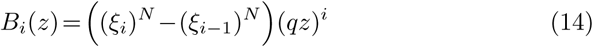

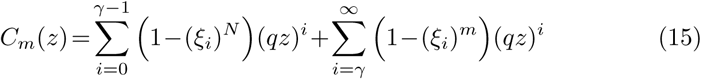

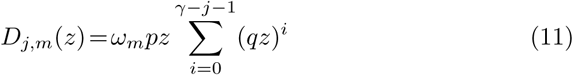

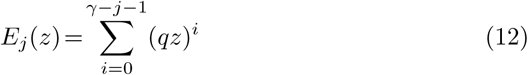

and where

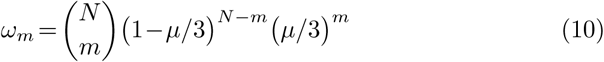

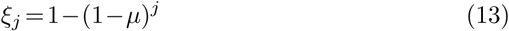

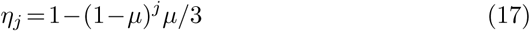

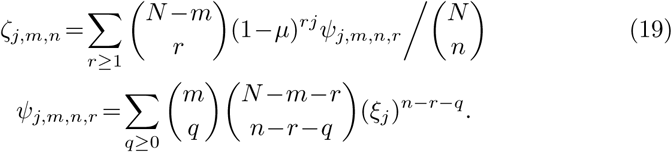

***Remark 4.** In the special case N = 0, the transfer matrix simplifies to the extent that we can compute the weighted generating function of the reads without on-target MEM seed in closed form. The result is*

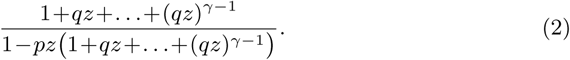

*Expression (2) was shown in section 3.4 to be the weighted generating function of reads without on-target exact seed. This shows that when there are no duplicates, MEM seeds have exactly the same properties as exact seeds*.

### 6.6 Computing MEM seeding probabilities

The matrix 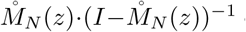 contains the weighted generating functions of all the reads without on-target MEM seeds. The term of interest, as usual, is the top right entry. Indeed, every read can be prepended by ↓_/0_ segments and only by those, otherwise the read would start with fewer than *N* soft masks. Thus, reads without an on-target MEM seed are precisely the sequences of segments that can be appended to the symbol ↑_*/*0_ and that are terminated by a tail.

To compute this term, we proceed as in section 3.5, *i.e.*, we compute the powers of 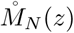 in the arithmetic of truncated polynomials and we stop the iterations when the terms are negligible. We bound the probability that the read contains more than *e* sequencing errors using the expression

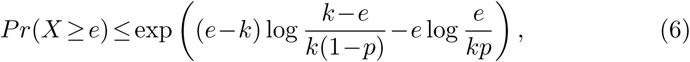

but here not every segment contains an error. There cannot be two symbols ↑_*/j*_ in a row, so a read with *s* +1 segments must contain a minimum number of sequencing errors which is *e* = └*s/*2┘. As before, we compute the powers of 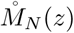 until the upper bound is less than a set fraction *ε* of the current value of *a*_*k*_.

If we call *M*_0_ the event that the read contains an on-target MEM seed, the method above gives us 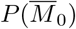. Calling *M*_*j*_ the event that the read contains a MEM seed for the *j*-th duplicate, we are interested in the probability

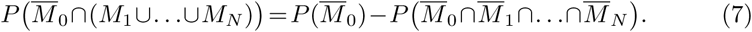

The key insight to compute 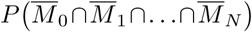 is to realize that there is some MEM seed, on-target or not, if and only if the read contains a match of size *γ* or more for any of the *N* +1 sequences. Therefore, this probability is the same as the term 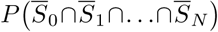 computed in section 4.

In conclusion, the probability that the MEM seeding process is off target is

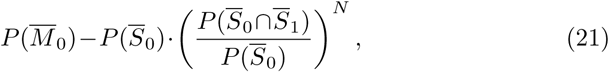

where 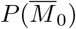 is computed using 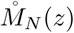 as explained in this section, 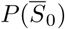 is computed using a recursive equation as explained in section 3.4, and 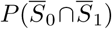 is computed using 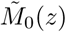 as explained in section 4.1.

### 6.7 Monte Carlo sampling

One potential difficulty in computing 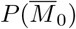 is that the matrix 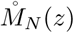 has dimension (*N* +*γ* +1)×(*N* +*γ* +1). The problem can become computationally intractable because *N* can be very large. For instance, the sequences called *Alu* have more than one million duplicates in the human genome. There is no hope to compute the powers of 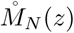 in these conditions and we need an alternative method.

The symbolic representation as MEM segments can be used to design an efficient method to sample reads. Instead of generating the nucleotides of the *N* + 1 sequences one by one, one can generate a single sequence of segments. Since the number of segment does not depend on *N*, we can obtain a fast Monte Carlo method to sample millions of reads and count the proportion that contain an on-target MEM seed.

The principle is to proceed in cycles of two steps. We first sample the position of the next sequencing error, which gives the position of the next symbol ↓_*/m*_, where *m* will be determined at a later stage. The second step is to determine whether there is a symbol *↑*_/*j*_ before that. For this we sample the number of masks that vanish before the symbol ↓_*/m*_. If they all vanish, the read contains an on-target MEM seed, provided the next read error is at a distance greater than *γ*. Otherwise, we sample the number *m* of hard masks at the sequencing error, and the process is repeated until we generate an on-target MEM seed, or until the read has size *k* or greater (in which case it has no on-target MEM seed).

The method is summarized in algorithm 1 below. It requires efficient algorithms to sample from the geometric and from the binomial distributions. Sampling from a geometric distribution can be done by computing the logarithm of a uniform (0, 1) random variable. Sampling from a binomial distribution can be done by the method of Kachitvichyanukul and Schmeiser [35]. Most importantly, the number of duplicates *N* has little influence on the running speed of algorithm 1.

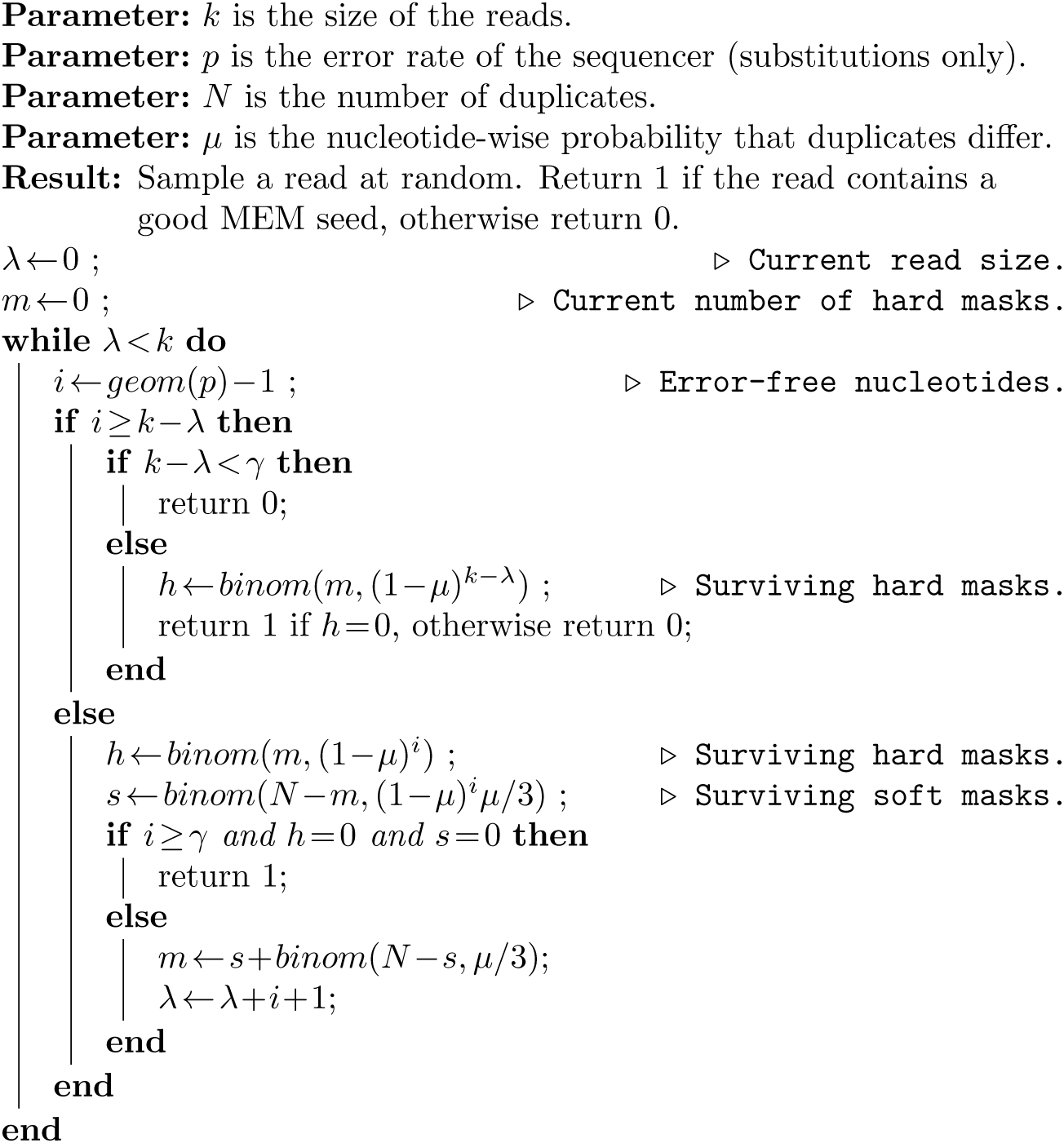

Algorithm 1 is an important result. It gives a compact solution to the problem of estimating the probability that a read can be mapped when using MEM seeds. The algorithm is also much faster than the naive approach of sampling every nucleotide of every sequence, because it is equivalent to sampling the nucleotide sequence of *all* the duplicates.

### 6.8 Illustration

We illustrate the strategy delineated above using the same settings as in section 4.2 (*k* =50, *p* =0.01 and *γ* =17), except that we replace exact seed by MEM seeds.

Fig. 14 shows the result for a number of duplicates *N* from 1 to 10 and for a divergence rate *µ* from 0 to 0.20. The curves have the same general aspect as those of Fig. 9. The probability that seeding is off target increases with *N*, as shown by expression (21).

**Figure 14:**
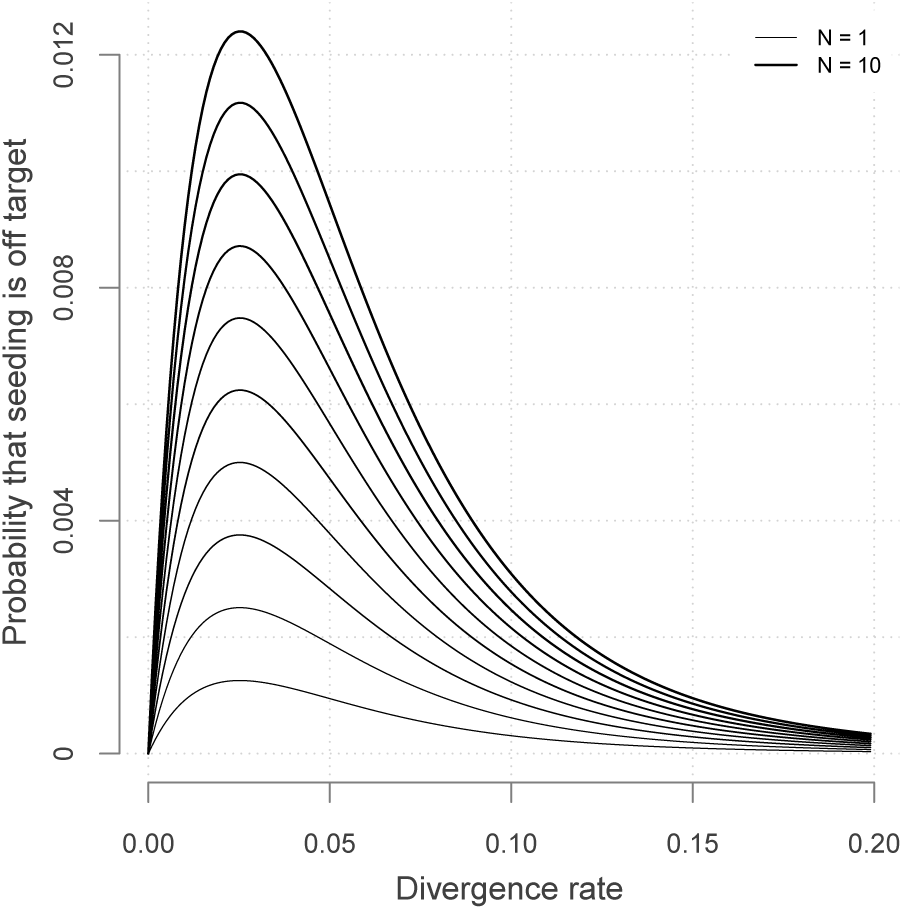
Off target seeding probabilities (MEM seeds). The curves show the probability that seeding is off target for MEM seeds of minimum size *γ* = 17. Read size and error rate are the same as in Fig. 9, *i.e. k* = 50 and *p* = 0.01. Each line shows a different number of duplicate sequences *N* from 1 to 10 and the x-axis shows the divergence rate *µ*, defined as the probability that a given duplicate differs from the target at any given position.

There is again a worst value of *µ* but it is much lower than the previous two cases (here it is close to 0.026). In the case of MEM seeds, it is not obvious why the same value of *µ* maximizes (21) for all values of *N* because both 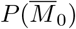 and 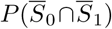 depend on *µ*.

Fig. 14 reveals that in this concrete case, MEM seeds increase the chances that seeding is off target by a factor up to 40 compared to exact seeds (see Fig. 9). On this criterion, MEM seeds are always inferior to exact seeds with the same specifications. The reason is that every read triggering an off-target seeding with exact seeds will also trigger an off-target seeding with MEM seeds. Knowing that, it is still important to realize how large the increase is in practice. MEM seeds tremendously simplify the seeding process, but this comes at the cost of an increase in the false positive rate. The methods presented here are thus very useful to balance the risk when using MEM seeds.

## 7 Random seeds

We have assumed throughout that seeds can match only the target or one of its duplicates. In practice, seeds can match many random locations of the genome and not just the target and the duplicates.

It is important for the validity of the theory that all the sequences that have a seed are considered candidate locations (as long as the seed is above the threshold *γ*). If not, the final estimates are biased. The hope is that spurious matches are shorter than *γ* so that they are filtered out, but this is not always the case. And if they qualify as a seed we need to verify the candidate location.

Such random hits can be problematic for two reasons: First, the time spent verifying them is wasted. Second, they can cause false positives. Fortunately, both issues can be addressed.

To save time during the alignment phase, we can prioritize the seeds so that the best hit is likely to be discovered first, allowing us to bail out from other alignments as early as possible. We can even bound the maximum quality of some candidates so that the alignment can be skipped altogether. These considerations depend on the implementation of the mapper so we will not develop this further. What matters is that random hits do not impose a significant burden on the time needed to verify the candidates.

The second issue is that random hits can generate false positives. Such cases occur when seeding is null (meaning that there is no seed for the target nor for its duplicates). Since the hits are not homologous to the read, the alignment score is extreme, which makes it easy to detect.

Indeed, if the candidate location is random, mismatches occur with probability 3/4. If it is the true location, mismatches occur with probability *p*. We discard the seed because it is an automatic match of size *γ* (and possibly larger in the case of MEM seeds). Say that there remain *L* nucleotides and that *m* of them are mismatches for the candidate location. From Bayes formula, the probability that the location is random given the number of mismatches is

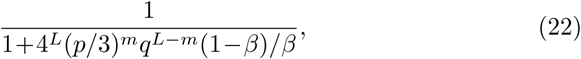

where *q* = 1 − *p* and *β* is the prior probability that the hit is random.

The value of *β* has little importance if *p* is small. Say that *p* = 0.01 and that a read generates a hit in the genome such that 33 nucleotides need to be aligned after seeding. If the hit is random there is a 99.99% chance that at least 15 are mismatches. The denominator of expression (22) is approximately 1+4.3⋅10^*−*18^(1 − *β*)*/β*. So unless *β <* 10^*−*17^, the support for the hypothesis that the hit is random is overwhelming. Conversely, if the hit is not random, there is a 99.99% chance that 4 or fewer nucleotides are mismathes. In this case, the denominator is approximately 1+6.8⋅10^9^(1 − *β*)*/β*, so unless 1 − *β <* 10^*−*9^, the support for the hypothesis that the hit is not random is overwhelming. If *p* is small, we do not need to worry about the value of *β*, one can choose for instance *β* = 1*/*2 so that the term (1*−β*)*/β* disappears from expression (22).

Since the probability that the best candidate is a random sequence is either very small or very large, it has no influence in the first case, or it dominates the probability of a false positive in the secondi case. For every read, we can use as confidence score the maximum of this probability and of the estimated probability that seeding is off target.

In summary, seeds in random sequences happen relatively frequently, but it is possible to minimize their computational burden. Also, they are no cause for concern regarding false positives because they can easily be detected using Bayes formula, as shown in expression (22).

## 8 The Sesame library

We implemented the methods and algorithms presented here in an open-source C library to compute seeding probabilities. The library is called Sesame and is available at https://github.com/gui11aume/sesame.

### 8.1 Main features of Sesame

Sesame contains functions to compute the seeding probabilities described here. All the functions were tested against simulations to ensure that the implementation is accurate and the code was checked extensively by static analysis and unit testing.

Computating seeding probabilities can take up to a few seconds, even when replacing iterative methods by Monte Carlo simulations. This is incompatible with the speed requirements of modern mappers, so Sesame has an interface for computations that must be performed online. In this mode, Sesame uses a type of lazy evaluation where the results are computed only the first time, and stored in memory for reuse on subsequent calls.

Storing the results in memory is an efficient strategy because three parameters are constant throughout the sequencing run: the minimum seed size *γ*, the read size *k* and the error rate of the sequencer *p* (and when using skip seeds, the amount of skipping *n* is also constant). Only two parameters depend on the read: the number of duplicates *N* and their divergence rate *µ*. In this mode, Sesame automatically switches to Monte Carlo sampling when *N* is large to save time. Also, the input parameters are “snapped” to a predefined grid of set values for *N* and *µ*, so that few computations are performed and most of the calls are actually memory lookups. Sesame can thus be integrated in short read mappers without being a bottleneck.

Alternatively, the probabilities of interest can be computed offline, saved to disk and loaded at run time. This is particularly useful if the sequencing runs follow some standard conditions with a known error rate, because the computations can be recycled between runs.

Finally, Sesame also has an offline interface, where seeding probabilities are computed exactly as requested by the users, *i.e.* without modifying the algorithm or the parameters, and also without storing the results in memory.

The Sesame manual, available from the repository, contains additional information and explains in detail how to use the library.

### 8.2 Using Sesame to compare seeding strategies

As a tool to compute seeding probabilities, Sesame can be used to compare the merits of different strategies. The kind of insight that we can gain from such calculations was already showcased in Fig. 9, Fig. 11 and Fig. 14, where the numbers were computed using Sesame.

To further showcase the potential benefits of computing seeding probabilities, we use Sesame to compare the default seeding strategies of BWA-MEM [30] and Bowtie2 [29]. Note that both mappers use advanced techniques to refine the seeds, so this comparison does not reflect the true performance of the mappers. It is nevertheless useful to know the baseline of each strategy. The default of BWA-MEM it to use MEM seeds of minimum size 19; that of Bowtie2 is to use skip-9 seeds of size 16.

The probabilities that seeding is off target for different read sizes *k* and different number of duplicates *N* are plotted in Fig. 15. The left panel shows the results for MEM seeds and the right panel shows the results for skip seeds. Here the error rate *p* is set to 1%, close to the specifications the Illumina platform, and the divergence rate between duplicates *µ* is set to an arbitrary value of 6%.

**Figure 15:**
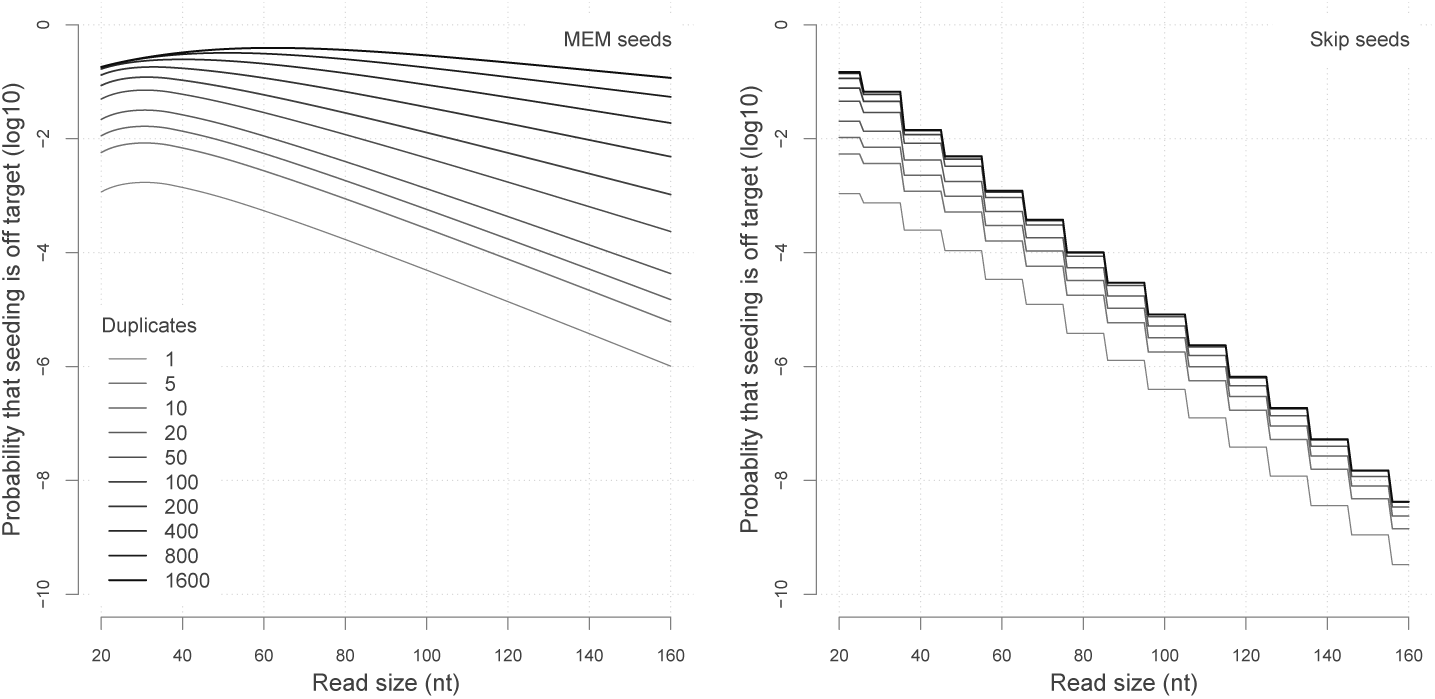
Comparing seeding strategies with Sesame. Left: MEM seeds with *γ* = 19 and *µ* = 0.06. Right: skip-9 seeds with *γ* = 16 and *µ* = 0.06. The probabilities that seeding is off target were computed using Sesame. Each curve represents the probability for a given number of duplicates (*N*). Estimates using iterative methods (skip seeds and MEM sees where *N ≤* 20) were computed to within 1% accuracy. Estimates using Monte Carlo sampling (MEM seeds where *N >* 20) were computed as the average of 500 million simulations.

The behaviors of the two types of seeds are dramatically different. Let us start with MEM seeds. For every value of *N*, the probability initially increases with the read size, and then drops exponentially. The initial increase is a hallmark of MEM seeds; it is due to the fact that duplicates can mask the target. Note that the asymptotic decay depends on the number of duplicates *N* because each duplicate can mask the target and thereby reduces the probability that it is discovered. Overall, these results show that the performance of MEM seeds is poor when the target has more than approximately 20 duplicates.

Turning to skip seeds, we see that the curves have a staircase look with a drop every 10 nucleotides. This is so because seeding probabilities remain unchanged until there is space for another seed of size 16 on the read. The curves are otherwise decreasing with a steady exponential trend where the asymptotic decay does not depend on the number of duplicates *N*. The reason is that duplicates do not prevent the target from being discovered, they merely fool the mapper when the target was not found. This property makes the asymptotic decay of skip seeds substantially faster that of MEM seeds when *N* is large.

Those properties are self evident in retrospect, but they are not necessarily obvious from the definitions of MEM seeds and skip seeds. The main limitations of the MEM seeds are best understood by keeping in mind that their asymptotic decay bepends on *N*.

Fig. 15 suggests that skip-9 seeds of size 16 are just better than MEM seeds of size 19. However, the gain in sensitivity comes at the cost of a larger candidate set, slowing down the mapping process. To remain competitive, Bowtie2 further filters the candidate set using a priority policy. But since some candidates are not checked, the probability that seeding is off target is larger than shown in Figure 15.

### 8.3 Key insights about MEM seeds

At least two key insights about MEM seeds can be gained from Fig. 15. The first is that it is probably not worth for a MEM-based mapper to check all the candidate loci when there are more than approximately 20 them. The mapper may find the correct location, but even if this is the case, the mapping quality will remain low because the prior chances of failure were high. A better strategy is to either bail out ot not waste time or to switch to a more sensitive seeding method (BWA opts for the second and uses a re-seeding policy). It is also important to note that this decision should be based on an estimated value of *N* and not, for instance, on the size of the seed or some other variable.

A second insight is that for MEM seeds, the off-target rate is always above 10^*−*3^ for reads of 50 nucleotides or fewer. Here it is important to mention that the value *µ* = 0.06 is not even the worst for reads of this size when *p* = 0.01 (according to Fig. 14, the worst value is around 0.02). So if *µ* is unknown and one wishes to be conservative, it seems that reads of 50 nucleotides cannot be mapped with confidence better than 1*/*1000.

However, this is not true for the reason that it is practically impossible to map an Illumina read of size 50 to the wrong location when *N* =0 (see section 7). This means that tweaking the seeding method to improve the mapping quality score will have less impact than checking whether *N* is 0 (in other words, whether the target is a unique sequence).

These insights suggest that there is a way to make MEM seeds more useful for short read mappers. We are currently working on a prototype based on these two ideas to increase the speed and the accuracy of short read mapping. This requires estimating *N* and *µ* for every read and will be described in a separate work.

## 9 Discussion

We have devised a set of methods to compute the probability that seeding heuristics fail and commit the mapping process to an error. We have also implemented the solution as an open source C library to perform the computations. This fills a knowledge gap to understand and calibrate the performance of the seeding heuristics. The pillars of our strategy are borrowed from analytic combinatorics [32, 33], even though we do not follow the complete programme. Constructing generating functions usually serves the purpose of finding their singularities in order to approximate the solution. In our case, however, the weighted generating functions cannot be computed so this strategy is not applicable.

To find the probabilities of interest in the absence of a fully specified weighted generating function, we compute only the first terms using iterative methods as explained in section 3.5, or using Monte Carlo sampling as explained in section 6.7. In this regard, the breakthrough is the encoding of reads as segments in different alphabets, which is an implicit form of Markov imbedding [36].

The methods rely on the knowledge of two essential parameters: the number of duplicates *N* and their divergence rate *µ*. These quantities can be estimated efficiently using the FM-index [21] and we will publish the detail of this method elsewhere, together with a more complete overview of how to use the results presented here to implement practical mapping algorithms.

The method is general, but it is important to clearly state the assumptions it depends on. First and most importantly, we have ignored insertions and deletions. We assume that the sequencing errors are substitutions only, which makes the method adapted to the Illumina technology, but not to deletion-prone instruments such as the Oxford Nanopore technology. We also assume that insertions and deletions never occur among duplicated sequences. This is obviously incorrect, but our initial tests with real data suggest that this is a minor impediment (most likely because this is only one of several simplifying assumptions). Incorporating insertions and deletions would make the theory intractable, so it is presently unclear how to deal with this type of error.

The second assumption is that all the candidate locations with a seed are tested with an exact sequence alignment algorithm, so that the best location is always discovered if it is present in the candidate set. This is not completely unrealistic, but it is important to note that for plant and animal genomes, a single seed may have tens of thousands of hits. Therefore, most mappers impose a limit on the number of alignments per read, opening the possibility that the best hit is seeded but not aligned. This is again a minor impediment, since we can replace the probability that the location is false by the proportion of candidates that were not aligned.

The third assumption is that all the duplicate sequences evolve independently of each other and at the same rate. This is again incorrect, because duplication events can happen continuously, creating complex ancestry relationships. It is possible to infer the ancestry using tree reconstruction techniques, but it would be challenging to incorporate this information in the present theory. The symbols of the alphabets developed above implicitly assume that the sequences are exchangeable and the complexity of the calculations explodes if it is not the case.

The last assumption is that seeds can match only the target or one of its duplicates. This does not hold in general but we have shown how to deal with this possibility *a posteriori* in section 7.

Being able to compute seeding probabilities revealed some interesting facts (sections 4.2, 5.2 and 6.8). The first is that the seeding schemes considered here have a worst case scenario for a particular value of the divergence between duplicates *µ*. Importantly, the worst value varies between different seeding methods, so it is possible in theory to construct opportunistic seeding strategies that pick the best method for every read.

Another interesting fact is that skip seeds can have better performance than exact seeds in the sense that they can yield lower off-seeding probabilities. However, this always comes at the cost of accuracy because skipping nulceotides reduces the probability of on-target seeding.

Finally, we also observed that MEM seeds cause a very serious increase in off-target seeding compared to exact seeds and skip seeds. This does not mean that MEM seeding is a bad strategy (it is usually faster than the other methods), but it is a good idea to keep an eye on performance and switch method or even skip the read altogether if the chances of discovering the target are too low.

Regarding the methodology, the Monte Carlo approach of algorithm 1 is relatively straightforward, so one may wonder why the approach with weighted generating functions is necessary for MEM seeds. The only reason is precision. To estimate the frequency of an event by Monte Carlo sampling, it must occur at least a few times in the simulation. For instance, with 1 million rounds of sampling, frequencies around 1*/*100, 000 or lower cannot be measured accurately. When one is interested only in frequent events, it is thus a reasonable strategy. On the other hand, for *N <* 20, the probability that MEM seeding is null or off-target is relatively small, so we need a method that is accurate in this range. Fortunately, the transfer matrix method is fast because the dimension of the matrix 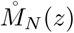 is small and the computations are not prohibitive.

The proposed methods meet the demand for speed. One needs to compute the probabilities only once for a given value of *N* and *µ* (the error rate *p* is known and constant). For *N >* 20, the iterative method is usually too slow and we need to use Monte Carlo sampling instead. The running time depends on *p*, on the size of the reads *k*, and on the desired number of iterations. Since those are constant throughout the sequencing run, the method always takes the same time (around 1-10 seconds for reads of size 100-200 nucleotides on modern hardware). The values of *N* and *µ* can be binned in intervals so that there are only around 100 pairs for a total cost of a few minutes per run. Considering that mappers can process at most 10, 000 reads per second per core, the time of mapping a sequencing run of 250 million reads is over 7 hours per core, two orders of magnitude larger than the time required to estimate the probabilities of error.

Finally, one may wonder if our approach has any advantage over methods based on machine learning. Such algorithms have already proved useful [37] and the rapid progress in the field of deep learning suggests that it is possible to train algorithms to accurately estimate mapping quality. In time, such algorithms may prove faster and/or more robust because they could learn intrinsic biases of the mapping algorithms. Yet, the main benefit of our approach will remain: the combinatorial constructions are a direct access to the nature of the problem. For instance, viewing MEM seeds through the lens of hard and soft masks turns a seemingly intractable process into a relatively simple one (see algorithm 1). The combinatorial stance is that there is value in clarity.

In conclusion, we presented a practical solution to the problem of estimating the probability of false positives when using seeding heuristics. This solution is adapted for mapping short reads sequenced with the Illumina technology. Being able to calibrate the seeding heuristic not only allows the user to choose how to balance speed versus accuracy, but also opens new applications. For instance, one can map reads from contaminated samples in pools of closely related genomes (*e.g.*, modern human and Neanderthal) in order to assign the reads to the organism they belong to. In this case, the probabilities of false positives give the right level of confidence in the assignment.

More generally, the analytic combinatorics programme is a very powerful tool to address problems in bioinformatics. Here we have seen how this strategy can be useful even when the generating functions cannot be computed. Using the same ideas, one could calibrate heuristics used in other alignment methods, especially in the expanding field of long-read technologies.

## Acknowledgements

We acknowledge the financial support of the Spanish Ministry of Economy, Industry and Competitiveness (‘Centro de Excelencia Severo Ochoa 2013-2017’, Plan Estatal PGC2018-099807-B-I00), of the CERCA Programme / Generalitat de Catalunya, and of the European Research Council (Synergy Grant 609989). R. C. was supported by the People Programme (Marie Curie Actions) of the European Union’s Seventh Framework Programme (FP7/2007-2013) under REA grant agreement 608959. We also acknowledge support of the Spanish Ministry of Economy and Competitiveness (MEIC) to the EMBL partnership.

